# Decreased ventral tegmental area CB1R signaling reduces sign-tracking and shifts cue-outcome dynamics in rat nucleus accumbens

**DOI:** 10.1101/2022.07.22.501038

**Authors:** Sam Z. Bacharach, David A. Martin, Cassie A. Stapf, Fangmiao Sun, Yulong Li, Joseph F. Cheer, Donna J. Calu

## Abstract

Sign-tracking rats show enhanced cue sensitivity before drug experience that predicts greater discrete cue-induced drug-seeking compared to goal-tracking or intermediate-rats. Cue-evoked dopamine in the nucleus Accumbens (NAc) is a neurobiological signature of sign-tracking behaviors. Here, we examine a critical regulator of the dopamine system; endocannabinoids, which bind the cannabinoid receptor-1 (CB1R) in the Ventral Tegmental Area (VTA) to control cue-evoked striatal dopamine levels. We use cell-type specific optogenetics, intra-VTA pharmacology and fiber photometry to test the hypothesis that VTA CB1R receptor signaling regulates NAc dopamine levels to control sign-tracking. We trained rats in a Pavlovian lever autoshaping task (PLA) to determine their tracking groups before testing the effect of VTA→NAc dopamine inhibition. We found this circuit is critical for mediating the vigor of the ST response. Upstream of this circuit, intra-VTA infusions of rimonabant, a CB1R inverse agonist, during PLA decrease lever and increase foodcup approach in sign-trackers. Using fiber photometry to measure fluorescent signals from dopamine sensor, GRAB_DA_, we tested the effects of intra-VTA rimonabant on NAc dopamine dynamics during autoshaping. We found that intra-VTA rimonabant decreased sign-tracking behaviors, which was associated with increases NAc shell, but not core, dopamine levels during reward delivery (US). We also observed a relationship between cue (CS)-evoked NAc dopamine activity and rigidity of behavior between rimonabant treatment sessions. Our results suggest that CB1R signaling in the VTA influences the balance between the CS- and US-evoked dopamine responses in the NAc and biases behavioral responding to cues in sign-tracking rats.

**SIGNIFICANCE STATEMENT:** Substance Use Disorder is a chronically relapsing neurobiological disorder that affects a subset of individuals that engage in drug use. Recent research suggests that there are individual behavioral and neurobiological differences prior to drug experience that predict addiction and relapse vulnerabilities. Here, we investigate how midbrain endocannabinoids regulate a brain pathway that is exclusively involved in driving cue-motivated behaviors of sign-tracking rats. This work contributes to our mechanistic understanding of individual vulnerabilities to cue-triggered natural reward seeking that have relevance for drug motivated behaviors.

## INTRODUCTION

Cues that reliably predict outcomes in the environment powerfully regulate behavior across conditioning. In addition to their predictive value, some cues gain enhanced motivational value that can lead to, or be associated with, maladaptive behavior. Training in a Pavlovian lever autoshaping (PLA) task reveals distinct conditioned responding phenotypes: Sign-tracking (ST) rats that engage predominantly with an insertable lever cue show enhanced discrete-cue induced relapse after cocaine experience. This stands in contrast to goal-tracking (GT) and intermediate (INT) rats that engage more with the foodcup and show lower levels of discrete-cue induced relapse after cocaine experience (Hearst & Jenkins, 1974; Tomie, 1996; Flagel et al., 2007; Meyer et al., 2012). Prior studies establish a role for NAc dopamine and endocannabinoid signaling in driving sign-tracking (Flagel et al., 2011; Bacharach et al., 2018). Here we investigate the extent to which midbrain endocannabinoid signaling and downstream Nucleus Accumbens dopamine (DA) dynamics interact to drive sign-tracking behaviors.

Cue-evoked dopamine release in the nucleus Accumbens (NAc) core distinguishes ST from GT phenotypes (Flagel et al., 2011). While pharmacological studies indicate that dopamine signaling drives the acquisition and expression of both sign- and goal-tracking behaviors (Danna & Elmer, 2010; Lopez et al., 2015; Fraser et al., 2016), sub-second cue-evoked NAc core dopamine is strongly correlated with and necessary for sign-tracking, but not goal-tracking behavior (Flagel et al., 2011; Saunders & Robinson, 2012; Clark et al., 2013; Fraser & Janak, 2017). Uncovering factors controlling this tracking-related difference in striatal dopamine signaling is crucial to understand the neurobiological mechanisms driving individual differences in motivation towards reward predictive cues.

Because of their role in regulating the dopamine system, we hypothesize that midbrain endocannabinoid signaling drives differences in cue-evoked striatal dopamine in sign- and goal-tracking phenotypes. Endocannabinoids are critical regulators of the dopamine system and blocking CB1R decreases striatal dopamine release (Cheer et al., 2000; Lupica et al., 2004; Cheer et al., 2004; Lupica & Riegel, 2005; Cheer et al., 2007; Oleson et al., 2012; Wenzel et al., 2018). The Ventral Tegmental Area (VTA) provides dense dopaminergic projections to the NAc, which are regulated by endocannabinoids (eCBs) acting at pre-synaptic CB1R to influence dopamine neuron firing and striatal dopamine release (Cheer et al., 2004; Lupica & Riegel, 2005). Blocking CB1R signaling decreases natural and drug reward self-administration, cue-induced reinstatement, and sign-tracking (McLaughlin et al., 2003; De Vries & Schoffelmeer, 2005; McLaughlin et al., 2006; Justinova et al., 2008; de Bruin et al., 2011; Oleson et al., 2012; Schindler et al., 2016; Bacharach et al., 2018). Blocking CB1R signaling in the VTA decreases both cue-evoked NAc dopamine and associated reward seeking (Oleson et al., 2012; Wenzel et al., 2018). We previously demonstrated that systemically blocking CB1R receptors decreases the enhanced attractive and reinforcing properties of lever cues in sign-tracking rats (Bacharach et al., 2018). We posit the locus of CB1R effects on sign-tracking is via VTA CB1R suppression of NAc dopamine release.

We hypothesize that CB1R receptor signaling in the VTA regulates cue-evoked NAc dopamine levels to control sign-tracking in rats. To test this, we use an optogenetic approach to examine the role of VTA→NAc dopamine projections in the expression of Pavlovian conditioned approach in ST, GT, and INT rats. We use intracranial pharmacology to examine the effect of intra-VTA infusion of the CB1R reverse agonist, rimonabant, on Pavlovian conditioned approach. Finally, combining intracranial pharmacology with fiber photometry we determine the extent to which VTA rimonabant affects cue- and reward-evoked NAc core and shell dopamine dynamics during Pavlovian conditioned approach. Altogether, we conclude from our data that CB1R signaling in the VTA maintains the conditioning-dependent behavioral and NAc DA bias towards cues in sign-tracking rats.

## MATERIALS AND METHODS

### Experimental subjects

We used female (n= 52) and male (n= 6) transgenic TH::Cre Sprague-Dawley rats (Envigo, Boyertown, PA; run in four cohorts) for the optogenetics experiment, female (n = 22) Long-Evans rats (Charles River Laboratories, Wilmington, MA; run in two cohorts) for the pharmacology experiment, and female (n=32) Long-Evans rats (Charles River Laboratories, Wilmington, MA; run in three cohorts) for the photometry experiments. All rats weighed between 215-350g at experimental onset. All rats were single-housed and maintained on a 12:12h light-dark cycle (ZT0 at 0730). All rats had ad libitum access to standard laboratory chow and tap water before food deprivation to 90% of their baseline weight, which was maintained for all experimental phases. Chow was provided after daily behavioral sessions. All procedures were performed in accordance with the “Guide for the care and use of laboratory animals” (8th edition, 2011, US National Research Council) and were approved by the University of Maryland School of Medicine Institutional Animal Care and Use Committee (IACUC).

### Surgical Procedures

For all surgeries, we anesthetized rats with 3 5% isoflurane (VetOne) and gave a subcutaneous injection of the analgesic, carprofen (5mg/kg). Prior to the first skull incision we gave rats a subdermal injection of 10 mg/mL lidocaine at the incision site. After lowering cannula or fiber optics into place, we secured them to the skull using jeweler’s screws and dental cement (Dentsply Caulk, Dentsply, York, PA). All coordinates given are distance from bregma according to the Paxinos and Watson rat brain atlas (6th edition, 2007).

#### Optogenetics

We infused 500n*L of* the Cre-dependent inhibitory chloride pump, halorhodopsin, AAV5-ef1a-DIO-eNphr3.0-eYFP (UNC vector Core, Chapel Hill, NC), or the control virus AAV5-ef1a-DIO-eYFP (UNC vector Core, Chapel Hill, NC) bilaterally in to the VTA (coordinates from bregma: -5.4 mm AP, ±2.15 mm ML, and −8.2 mm DV) at a rate of 100 nL/min using a microinfusion pump (UltraMicroPump III, World Precision Instruments, Sarasota, FL, USA) and a 10µL Hamilton syringe (Hamilton, Reno, NV, USA). The needle tip was left in place for 5 minutes after infusion, raised 0.1mm and left another 5 minutes before the final raising. We then implanted two 200 µm core, 0.67 NA fiber optics with ceramic zirconia ferrules (Prizmatix, Holon, Israel) targeting the NAc core, bilaterally (coordinates from bregma: +1.8 mm AP, +2.15 mm ML, -6.6 mm DV).

Fiber optics were secured with Denmat (Denmat, Lompoc, CA) then a thin layer of dental cement. For two of four cohorts we trained and tested rats in Pavlovian lever autoshaping six to eight weeks after surgery. In two of four cohorts, we gave rats (n = 14 Halo, n = 19 eYFP) three days of training before surgically injecting virus and implanting fiber optics, followed by a 6-week recovery/viral expression period, then a fourth training session before testing.

#### Pharmacology

We implanted guide cannulae (23G; Plastics One INC., Roanoke, VA) bilaterally into the VTA at a 10-degree angle (coordinates from bregma: -5.4 mm AP, ±2.2 mm ML, and −7.33 mm DV). Cannula were secured with jewelers screws and dental cement. At the end of surgery, we inserted dummy cannula into the guide cannula, which were kept in the guide cannula and were only removed during infusion habituation and infusion test procedures. Animals were given 2-3 weeks recovery before testing.

#### Fiber photometry

Guide cannulae were implanted in the VTA identical to *Pharmacology* experiments. In addition, we infused 1µL of AAV9-hSyn-DA2m (GRAB_DA_) into the Nucleus Accumbens unilaterally at a 6-degree angle (coordinates from bregma: +1.8 mm AP, +2.15 mm ML, -6.6 mm DV). Following virus infusion, we implanted one 400 µm core, 0.67 NA fiber optic (Thor labs) in the NAc, 0.1mm dorsal to the virus injection coordinates. Fiber optics were first cemented in place with Metabond (Parkell Inc., New York, New York), then with Denmat. The entire headcap was covered in a thin layer of dental cement. Rats were given 4 weeks for recovery before testing.

##### Histology

At the end of experiments, we deeply anesthetized rats with isoflurane and transcardially perfused them with 100 ml of 0.1M sodium phosphate buffer (PBS), followed by 400 ml of 4% paraformaldehyde (PFA) in PBS. We removed brains and postfixed them in 4% PFA for two hours before we transferred them to 30% sucrose in PBS for 48 -72h at 4 °C. We subsequently froze brains and stored them at −20 °C until sectioning. Coronal sections (50 μm) containing NAc and VTA were collected using a cryostat (Leica Microsystems).

To verify cannula placements, we stained brain sections with cresyl violet and coverslipped with Permount (Fischer Scientific, Waltham, MA). To verify viral expression and fiber placements we mounted and coverslipped all brain sections with Vectashield DAPI (Vector Laboratories, Burlingame, CA) and the mounting medium, Mowiol (Sigma-Aldrich, St. Louis, MO). We used a spinning disk confocal microscope (Leica SP8) to verify viral expression and fiber optic/cannula placement. For optogenetic experiments, rats were excluded from analysis if cell body labeling of Halo or eYFP was observed outside of the VTA (in SNc), if there was no terminal expression in the NAc core, and/or fiber optics were not targeting the NAc core. For pharmacology experiments rats were excluded if cannula were not targeting the VTA. For photometry experiments rats were excluded if there was no virus expression below the fiber optic in the NAc, or if the CS-evoked dopamine signal was less than 2 Zs above baseline during the Vehicle testing day.

##### Behavioral apparatus

Experiments were conducted in individual sound-isolated standard experimental chambers (25 × 27× 30cm; Med Associates). For Pavlovian lever autoshaping each chamber had one red house light (6 W) located at the top of a wall that was illuminated for the duration of each session. During PLA, the opposite wall in the chamber had a recessed food cup (with photo beam detectors) located 2 cm above the floor grid. The food cup was attached to a programmed pellet dispenser that delivered 45 mg food pellets (catalog#1811155; Test Diet 5TUL; protein 20.6%, fat 12.7%, carbohydrate 66.7%). One retractable lever was positioned on either side of the food cup, counterbalanced, 6cm above the floor.

#### Optogenetics

Each Med Associates chamber was equipped with a green LED (525 nm, max 300mW, Prizmatix) to deliver light to the NAc core. A TTL pulse was generated by MedPC 1s prior to lever insertion and was sent to an Arduino minicontroller (Somerville, MA) which drove an LED for 11s total, (terminating with the retraction of the lever). The complete light path is MED TTL→Arduino→LED→Patch cord (1000µm core, Prizmatix)→commutator (Prizmatix)→ bifurcated (2x500µm core, Prizmatix) fiber optic patch cord → 2 implanted fiber optics ferrules targeting the NAc core. A ceramic sleeve covered in black heat shrink tightly joined the patch cord and fiber optic and prevented light loss. We used a light meter (PM100D, Thorlabs) to calibrate light output in each box before and after each session.

#### Fiber photometry

We used LEDs (ThorLabs) to deliver 465 nm light to measure GRAB_DA_ fluorescence signals and 405 nm light as an isosbestic control. The two wavelengths of light were sinusoidally modulated at 210 and 337Hz respectively. The LEDs connected to a fluorescence mini cube (Doric Lenses). The combined LED output passed through a fiber optic cable (1 m long; 400 μm core; 0.48 NA; Doric) which was connected to implanted fiber optics (400 μm core, Thor Labs). We maintained the light intensity at the tip of the fiber optic cable at 10-15 µW across behavioral sessions. LED light collected from the GRAB_DA_ and isosbestic channels was focused onto a femtowatt photoreceiver (Newport). We low-pass filtered and digitized the emission light at 3Hz and 5 KHz respectively by a digital processor controlled by Synapse software suite (RZ5P, Tucker Davis Technologies (TDT)). We time-stamped the behavioral events including lever insertion, pellet delivery, lever press, and food cup entry through TTL pulses in Synapse software.

##### Training in Pavlovian lever autoshaping

Across all experiments, we gave rats a single 38-minute magazine training session during which one food pellet was delivered into the food cup on a variable interval (VI) 90 s schedule (60-120 s) for 25 trials. We trained rats in four or five daily Pavlovian lever autoshaping (PLA) sessions, which consisted of 25 reinforced lever conditioned stimulus (CS+) presentations occurring on a VI 90 s schedule (60-120 s). CS+ trials consisted of the insertion of a retractable lever for 10 s, after which the lever was retracted and two food pellets were delivered to the food cup regardless of whether a lever or food cup response was made. For two-lever PLA experiments, on the opposite side of the wall from the CS+, we included a CS- lever for which the extension and retraction had no programmed consequences.

Sessions consisted of 25 CS+ pairings and 25 CS- trials with a VI90s schedule between rewarded trials. For experiments in the unpaired condition, lever extension/retraction did not produce food pellets. Instead, two food pellets were delivered pseudorandomly during the ITI (at least 20s after/ 20s before lever retraction/extension, respectively).

#### Measurements and difference scores

Behavioral measurements were collected during the 20s pre-CS period (ITI), 10s CS period, and the 5s post-CS reward delivery period. An automated measurement of the latency to first contact the lever and/or food cup during the cue for each trial was recorded. On trials in which a contact did not occur, a latency of 10 s was recorded. For each session the lever or food cup probabilities were calculated by determining the number of trials that the lever or food cup response was made, divided by total number of trials in the session.

We used a Pavlovian Conditioned Approach (PavCA) analysis (Meyer et al., 2012) to determine sign-goal- and intermediate tracking groups. The PavCA score quantifies the difference between lever-directed and food cup-directed behaviors, and ranges from -1.0 to +1.0. An individual rat’s PavCA score is the average of three difference score measures (each ranging from -1.0 to 1.0) including: (1) preference score, (2) latency score and (3) probability score. The preference score is the total number of lever presses during the CS, minus the total number of food cup responses during the CS, divided by the sum of these two measures. The latency score is the session averaged latency to make a food cup response during the CS, minus the session averaged latency to lever press during the CS, divided by the duration of the CS (10s). The probability score is the probability of lever press minus the probability of food cup response observed across the session. ST PavCA scores range from +0.33 to +1.00, INT PavCA scores range from +0.32 to -0.32, and GT PavCA scores range from -0.33 to -1.00.

### Testing in Pavlovian lever autoshaping

#### Optogenetics

We habituated rats to the optogenetic patch cable after D4 of training. On subsequent sessions, rats were tethered to the patch cable during the session. During the OFF test, rats were tethered but no light was delivered. For the ON Test, light was delivered to the NAc at 4-6mW from the fiber tip. During the ON test, LED light delivery began 1s prior to lever extension and remained on for the duration of the 10 s lever cue (11s total). Lever retraction, termination of the LED light, and delivery of food occurred simultaneously at the end of each of the 25 trials.

#### Pharmacology

We habituated rats to handling and infusion procedures throughout training. After the last training day, we inserted injectors in to the cannulae and the infusion pump was turned on, but nothing was infused. Before test sessions in PLA, we gave each rat an infusion of rimonabant or vehicle in two separate counterbalanced test sessions that occurred 48 hours apart. We removed dummy cannulae and inserted 30G injector cannulae (Plastics One) extending 1.0 mm beyond the end of the guide cannulae. We connected each injector cannula using polyethylene-50 tubing, which was attached to a 5 μL Hamilton syringe (Hamilton, Reno, NV) that was placed in an infusion pump (CMA syringe pump 4004; Harvard apparatus). We infused rimonabant (2 µg/µL) or vehicle bilaterally into the VTA at a rate of 0.25 µL/min for a total of 2 minutes, or 0.5 µL total volume per hemisphere. We kept the injectors in place for an additional minute before slowly removing them and replacing dummy cannulae for behavioral testing. We tested rats in PLA 10 minutes after the completion of the infusion.

Drug solutions were prepared immediately before each test session. Rimonabant (SR141716A, 5-(4-Chlorophenyl)-1-(2,4-dichloro-phenyl)-4-methyl-N-(piperidin-1-yl)-1*H*-pyrazole-3-carboxamide, NIDA Drug Supply Program) was dissolved in a 1:1:18 solution of ethyl alcohol (Sigma), emulphor (Alkamuls EL-620, Solvay Chemicals, Princeton, NJ), and saline (Hospira) and sonicated for 15 min. The vehicle solution consisted of the 1:1:18 solution of ethyl alcohol, emulphor, and saline.

#### Photometry

We habituated rats to patch cables during magazine training and performed recordings during all of training and testing phases. Testing and drug infusions procedures were identical to pharmacology experiments.

##### Data and Statistical analyses

Behavioral data was analyzed using SPSS statistical software (IBM) with mixed-design repeated-measures ANOVA. Significant main and interaction effects (p < 0.05) were followed by post-hoc within-subject, repeated-measures ANOVA or t-tests. For significant post-hoc t-tests we report Cohen’s d effect size.

#### Optogenetics

For training data, we used mixed repeated measures ANOVA including within-subject factor of Session (4) and between-subject factors of Tracking (GT, INT, ST) and Virus (Halo, eYFP). For Test data, we used mixed repeated measures ANOVA including within-subject factor of Light (OFF, ON) and between-subject factors of Tracking (GT, INT, ST) and Virus (Halo, eYFP). In cases with a significant Treatment x Tracking interaction, we then probed individually Response x Treatment interactions within each tracking group.

#### Pharmacology

For PLA training data, we used mixed repeated measures ANOVA of lever and food cup measures (contact, latency and probability), using between-subject factors of Tracking group (ST, INT) and within-subject factor of Session to analyze lever- and food cup-directed behaviors. For Test data, we used mixed repeated measures ANOVA including within-subject factor of Treatment (Veh, Rimo) and between-subject factors of Tracking (INT, ST). In cases with a significant Treatment x Tracking interaction, we then probed individually Response x Treatment interactions within each tracking group.

#### Photometry

For training data, we used mixed repeated measures ANOVA including within-subject factor of Session (5). For Test data, we used mixed repeated measures ANOVA including within-subject factor of Treatment (Veh, Rimo) and Epoch (Cue, Reward).

#### Photometry signal analysis

To calculate ΔF/F, a least-squares linear fit was applied to the 405 nm signal. ΔF/F = (490 nm signal − fitted 405 nm signal)/fitted 405 nm signal. To calculate Z-scores on each trial, we took the average of the ΔF/F signal over a 10s baseline period before the CS insertion and divided the total trial measurements by that average. All 25 trials per session for each animal were averaged in to one average per animal, which was used for subsequent analysis. To screen for reliable photometry signal, we defined significant transients in a behavioral window (5s post lever insertion) as having a maximum CS-evoked peak amplitude ≥ 2 Z-score (*p*=0.05) above baseline. We excluded rats whose vehicle day CS-evoked signals did not meet this criterion. Peak height was calculated using the “findpeaks” function in Matlab. We took the maximum peak height above Z = 0 in the 2.5s following the event of interest (either CS (lever extension) or US (lever retraction and pellet delivery)). In the case there were no peaks above Z=0, the peak height measure was recorded as 0 for that subject. The latency at which the maximum peak occurred was used for the peak latency calculation. We used the “trapz” function in Matlab to examine area under the curve for 5s post event. For training data in **Fig. 4**, of n=22 rats, four were only recorded on D1 and D5. Thus, In **Fig. 4D inset** and **4F** data are from n=18 animals, while data in **4D main figure, 4E,** and **4G** are from n=22 animals. When examining differences in signal between NAc Core and Shell in figures **4H-J**, we analyze data from 20 of 22 rats because 2 fiber placements were exactly on core/shell border.

## RESULTS

### Individual differences in the acquisition of conditioned approach in a Pavlovian lever autoshaping task

We first examined the role of the VTA→NAc dopamine projection in regulating individual differences in Pavlovian approach. We infused a Cre-dependent control virus (AAV5-ef1a-DIO-eYFP) or a Cre-dependent Halorhodopsin (Halo) (AAV5-ef1a-DIO-eNphr3.0-eYFP) in to the VTA of TH::Cre rats and implanted bilateral optical fibers targeting the NAc core (**Fig. 1A, B**). After a recovery period, we trained rats (Halo, n = 31, eYFP, n = 27) in four Pavlovian lever autoshaping (PLA) sessions where the extension and retraction of a lever for 10s predicted the delivery of food (**Fig. 2A**). We classified rats as Sign-Trackers (ST), Goal-Trackers (GT), or Intermediates (INT) based on the Pavlovian conditioned approach (PavCA) score, which reflects an animals’ tendency to approach the lever relative to the food cup. Over acquisition we observed main effects of Session (F(3,156) = 34.921, *p* < 0.001) and Tracking (F(2,52) = 67.568, *p* < 0.001), and a Session x Tracking interaction (F(6,156) = 24.923, *p* < 0.001), confirming individual differences in conditioned approach (**Fig. 2B)**. Main effects and interactions of all behavioral measures collected in PLA are presented in **Table 1**. There were no main effects or interactions with Virus (Max F = 3.061, min *p* = 0.086), demonstrating that the virus infused had no effect on the acquisition of behaviors that characterize sign, goal- and intermediate rats.

**Figure 1:**
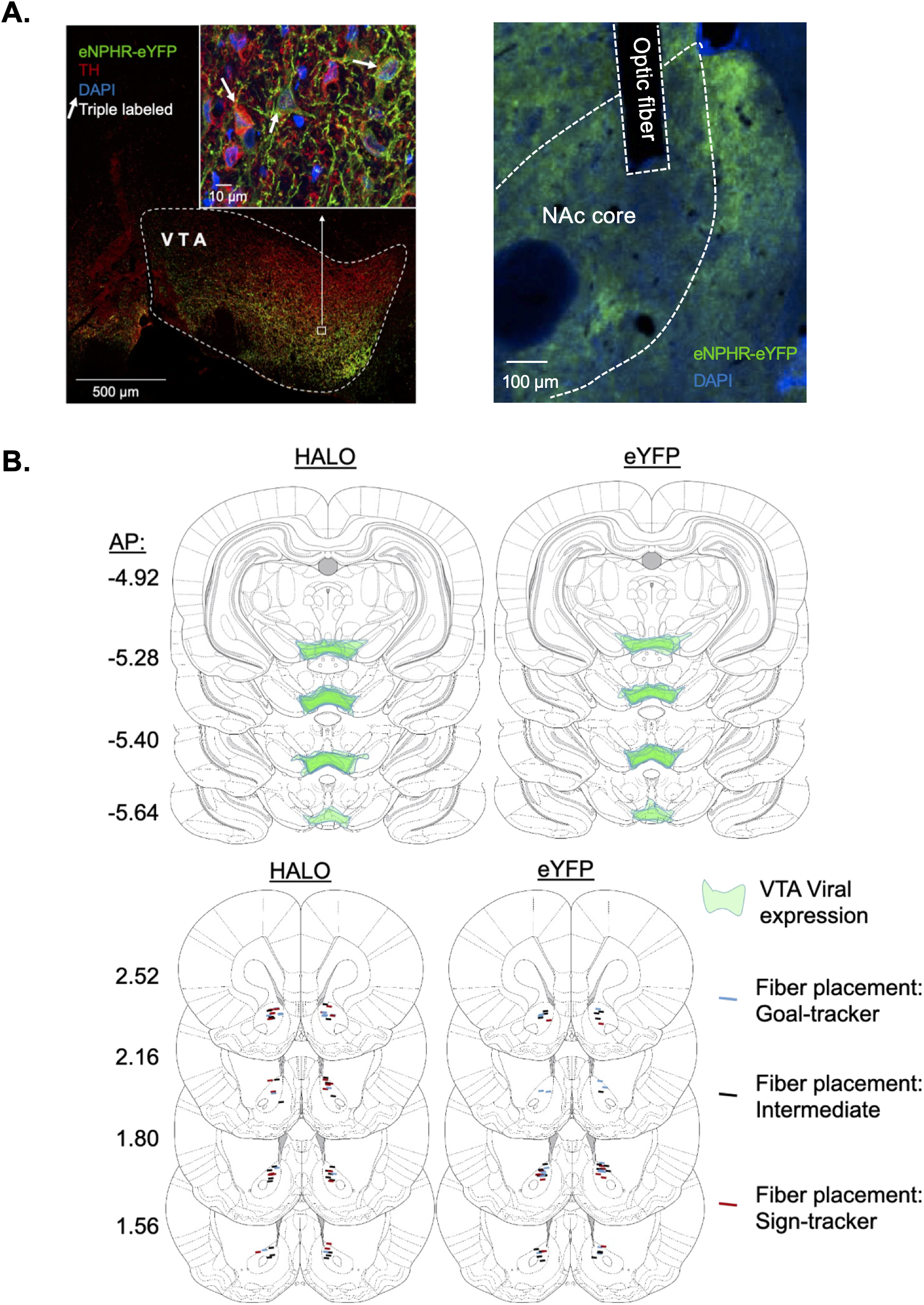
Histological verification of viral and optogenetic fiber placement. ***A***, (Left) Representative VTA transduction of Halorhodopsin (green). Staining for Tyrosine Hydryoxylase (TH) is given in red, DAPI in blue. White arrows depict triple overlap. (Right) Representative image Halorhodopsin terminal expression (green) and DAPI (blue) in the Nucleus Accumbens. White dotted line indicates boundary of NAc core. ***B***, (Top) The extent of viral coronal transduction (in mm) of Halorhodopsin and eYFP in the VTA. (Bottom) Bilateral optogenetic fiber optic placement in the NAc across the three tracking groups.

**Figure 2:**
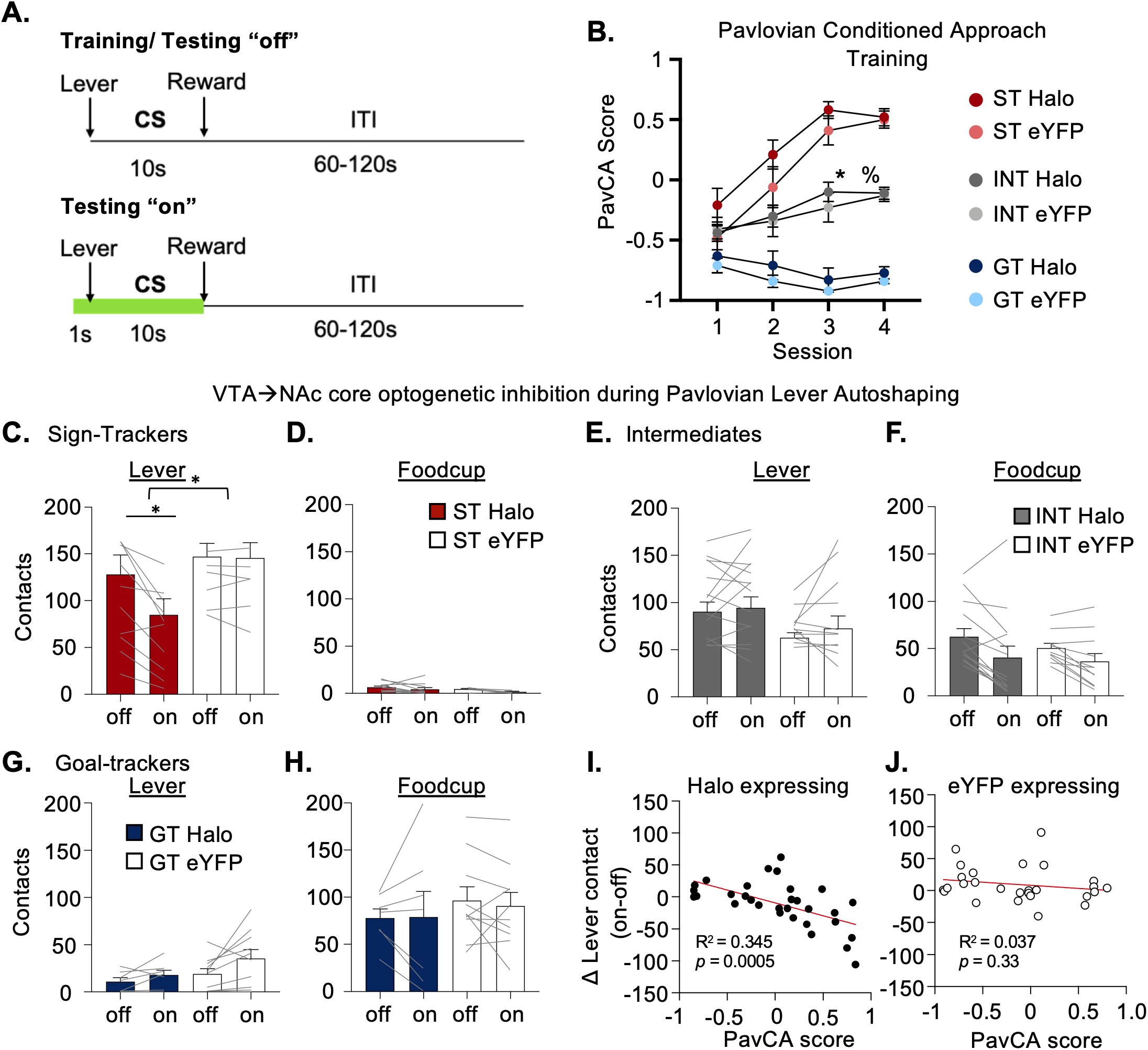
VTA-NAc dopamine terminal inhibition selectively reduces approach in sign-trackers. ***A***, After training in PLA, rats were given test sessions with light turned OFF(top) or ON(bottom) where light was delivered to the NAc core during the 10s cue period. ***B***, PavCA score mean ± SEM. ST, GT, and INT acquire individual differences in conditioned responding in PLA task. * Main effect of Session; % Significant Session x Tracking interaction. ***C***, Terminal inhibition of dopamine axons in the NAc significantly reduces the amount of lever contacts and ***D,*** has no effect on foodcup contacts in Sign-trackers. * With curved bracket indicates significant Light x Virus interaction. * With straight brackets indicates significant post-hoc within-subject t-test. ***E***, ***F***, No significant effects of terminal inhibition in INT rats. ***G***, ***H***, No significant effects of terminal inhibition in GT rats. ***I,*** Correlation between the change in lever pressing behavior between light conditions as a function of Halo-expressing rats PavCA scores. ***J,*** Correlation between the change in lever pressing behavior between light conditions and eYFP-expressing rats PavCA scores.

**Table 1:**

|  |  | Lever |  |  |  |  |  | Foodcup |  |  |  |  |  |
| --- | --- | --- | --- | --- | --- | --- | --- | --- | --- | --- | --- | --- | --- |
| Factor | Degrees Of Freedom | Contact |  | Latency |  | Probability |  | Contact |  | Latency |  | Probability |  |
|  |  | F | *p* | F | *p* | F | *p* | F | *p* | F | *p* | F | *p* |
| Session | (3,156) | 59.965 | <0.001 | 33.276 | <0.001 | 35.963 | <0.001 | 3.962 | 0.009 | 3.185 | 0.026 | 2.427 | 0.068 |
| Tracking | (2,52) | 43.241 | <0.001 | 38.632 | <0.001 | 80.645 | <0.001 | 12.628 | <0.001 | 22.588 | <0.001 | 25.657 | <0.001 |
| Virus | (1,52) | 1.537 | 0.221 | 0.125 | 0.726 | 0.875 | 0.354 | 0.383 | 0.539 | 3.198 | 0.08 | 3.831 | 0.056 |
| Session x Tracking | (6,156) | 23.015 | <0.001 | 10.522 | <0.001 | 10.116 | <0.001 | 14.473 | <0.001 | 13.420 | <0.01 | 13.967 | <0.01 |
| Session x Virus | (3,156) | 1.163 | 0.326 | 2.009 | 0.115 | 0.762 | 0.517 | 0.805 | 0.493 | 1.399 | 0.245 | 1.343 | 0.263 |
| Tracking x Virus | (2,52) | 0.299 | 0.743 | 0.236 | 0.791 | 0.182 | 0.834 | 1.081 | 0.347 | 1.694 | 0.194 | 2.306 | 0.110 |
| Session x Tracking x Virus | (6,156) | 0.887 | 0.506 | 1.070 | 0.383 | 1.078 | 0.378 | 0.791 | 0.578 | 0.484 | 0.820 | 0.470 | 0.830 |

### VTA-NAc dopamine terminal inhibition selectively reduces approach in sign-trackers

Prior pharmacology studies establish a role for NAc core dopamine in specifically driving sign-tracking (Clark et al., 2013; Fraser et al., 2016; Fraser & Janak, 2017; Saunders & Robinson, 2012). Here we test the role of VTA-NAc core projections by inhibiting VTA dopaminergic axons projecting to the NAc core during the 10s lever cue in sign-, goal- and intermediate tracking groups. We gave two test sessions; one in which the rat were tethered to the patch cable but light was not delivered (OFF condition) and a second session in which rats were tethered and LED light was delivered to the NAc core (ON condition) (**Fig. 2A**).

We examined the effect of dopamine inhibition on the number of lever contacts in the ST group (**Fig. 2C**) and found a main effect of Light (F(1,14) = 8.735, *p =* 0.010) and a Light x Virus interaction (F(1,14) = 67.602, *p =* 0.015). This interaction was driven by a decrease in pressing by the Halo group (*t*_(9)_ = 3.884, *p* = 0.004, Cohen’s d = 0.71) but not the eYFP group (*t*_(5)_ = 0.277, *p* = 0.793). We additionally examined foodcup contacts and found no significant main effects or interactions (Max F = 3.35, Min p = 0.089 (ME light)) (**Fig. 2D**). In addition to lever and foodcup contacts, we analyzed the latency and probability to lever and food cup contact and found no significant interactions with virus (Max F = 2.07, Min *p* = 0.172). Collectively, these results demonstrate that dopamine inhibition in ST rats doesn’t affect the latency or probability to make a response, but selectively reduces the vigor with which the animals engage with and press the lever. Consistent with pharmacological manipulations in the NAc core (Saunders & Robinson, 2012; Clark et al., 2013; Fraser & Janak, 2017), inhibition of the VTA→NAc dopamine projection is necessary for the vigor of lever responding in the sign-tracking rats.

We next examined the effect of dopamine terminal inhibition in INT rats. Even though the INT rats had substantial levels of lever pressing by the end of training, there was no Light x Virus interaction (F(1,23) = 0.208, *p* = 0.653) (**Fig. 2E**). Poking behavior remained unaffected by dopamine inhibition as well. Although there was a main effect of Light (F(1,23) = 19.504, *p <* 0.001), there was no Light x Virus interaction (F(1,14) = 0.848, *p =* 0.367) (**Fig. 2F**). Intermediate rats display similar amount of both lever and foodcup directed behaviors, and our results demonstrate that NAc dopamine inhibition does not affect the levels of either response. The preferred response of GT rats is foodcup directed behavior. There was no significant interaction with Light x Virus (F(1,15) = 0.125, *p =* 0.729) (**Fig. 2G**). We found a main effect of Light when examining pressing behavior (F(1,15) = 5.547, *p =* 0.033) (**Fig. 2H**), but no Light x Virus interaction (F(1,15) = 0.922, *p =* 0.352).

We performed an analysis of continuous data showing the relationship between PavCA score and the difference in lever contacts between treatments (ΔLever contacts = contacts light ON -contacts light OFF). We found a significant negative correlation (R^2^ = 0.3452, *p* = 0.0005 **Fig. 2I**), indicating the greater the PavCA score (ST) the larger the decrement in lever presses induced by dopamine inhibition. We did not see a significant correlation in the eYFP control group (R^2^ = 0.03746, *p* = 0.33, **Fig. 2J**), nor did we observe any relationship between PavCA score and difference in food cup responses between treatments (R^2^ = 0.0012, *p* = 0.8554; eYFP R^2^ = 0.0014, *p* = 0.853) (data not shown). To further confirm that inhibiting VTA dopaminergic axons in the NAc was specific to lever pressing of the ST, we analyzed data only from the rats expressing Halorhodopsin using within between-subjects factor of Tracking (ST, INT, GT) and within subjects factors of Light (OFF, ON) and Response (press, poke). We found a 3-way interaction (F(2,28) = 7.98, *p* = 0.002) that was driven by a decrease in pressing by the sign-tracking rats. This analysis further demonstrates that VTA dopamine axon inhibition specifically reduces lever pressing in sign-tracking rats.

In conclusion, inhibiting dopaminergic terminals in the NAc specifically reduced the amount of lever pressing in ST animals and had no effect on the responding in INT or GT rats. Here we have demonstrated this VTA→NAc pathway is important for regulating sign-tracking behavior, we next manipulate CB1R in the VTA where we predict CB1R regulates cue-evoked NAc DA levels during sign-tracking.

### Intra-VTA CB1R inhibition reduces sign-tracking

Given the specific effects of VTA→NAc core dopamine inhibition in sign-tracking rats, we next examine how VTA CB1R regulate cue attraction in sign-trackers. As a comparison group, we include INT rats who also display lever directed behaviors, but do not require NAc dopamine to execute these behaviors (**Fig. 2E,F)**. All rats were implanted with cannulae targeting the VTA (**Fig. 3A).** First, we trained 19 rats (11 ST, 8 INT) for 5 PLA sessions to determine tracking groups. For the PavCA score we observe main effects of Session (F(4,68) = 21.98, *p* < 0.001) and Tracking (F(1,17) = 22.77, *p* < 0.001) and a Session x Tracking interaction (F(4,68) = 3.45, *p* = 0.013) (**Fig. 3B**). On the fifth training session INT rats showed similar levels of lever directed and food cup directed behaviors (PavCA = 0.13) whereas ST rats showed predominantly lever-directed behaviors (PavCA = 0.62) further confirming these two groups of rats show different patterns of conditioned responding in response to Pavlovian reward-predictive cues by the end of training. We present main effects and interactions of all PLA training measures in **Table 2**.

**Figure 3:**
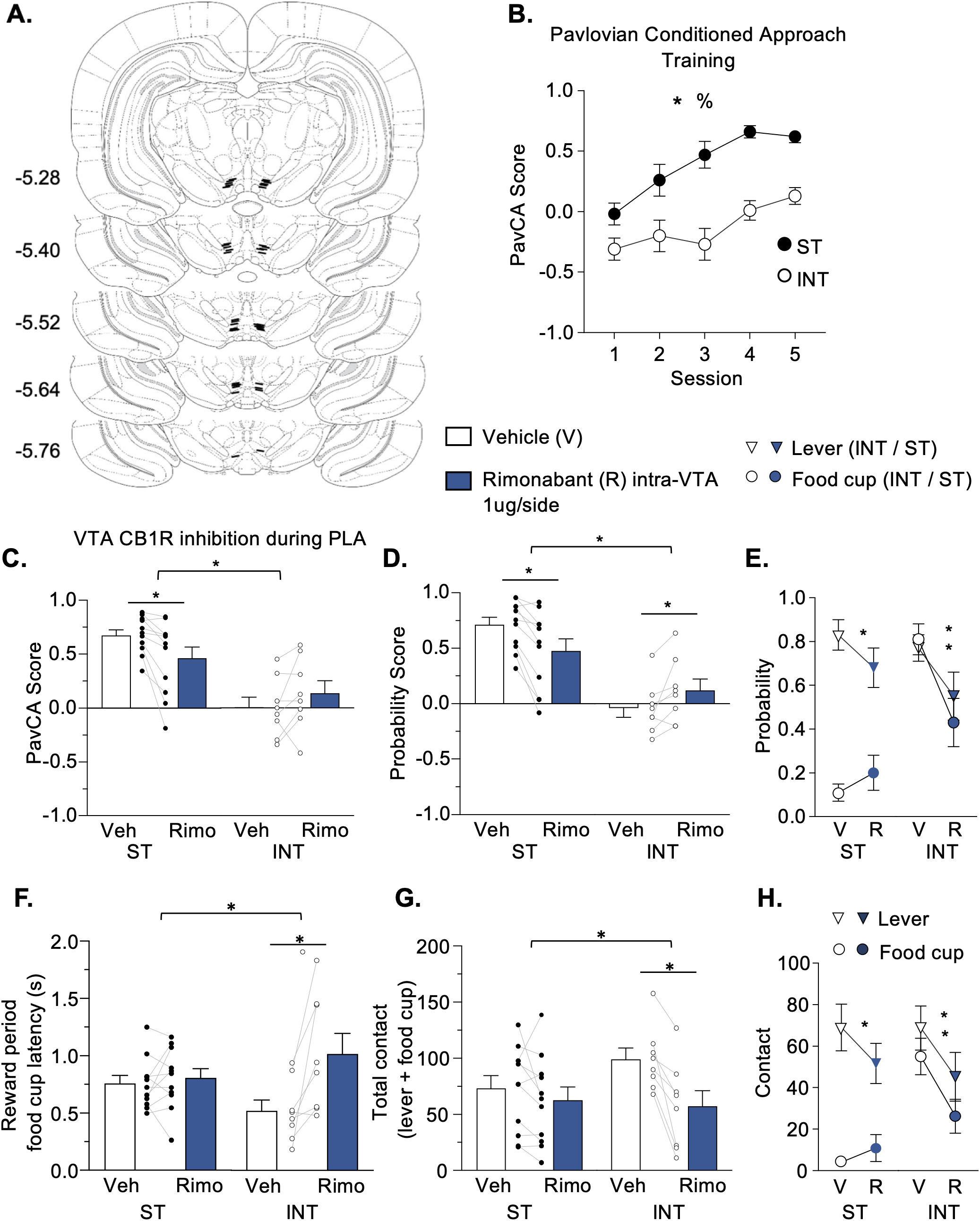
Intra-VTA CB1R receptor inhibition reduces sign-tracking. ***A***, Coronal sections (in mm) depicting location of VTA injector tips for rimonabant infusion. ***B***, PavCA score mean ± SEM of ST and INT that acquire individual differences in conditioned responding in PLA task. * Main effect of Session; % Significant Session x Tracking interaction. ***C***, Rimonabant significantly decreases PavCA in ST, but not in INT. * with curved bracket indicates significant Tracking x Treatment interaction. * With straight bracket indicates significant effect of treatment. **D**, ST show a significant reduction in the probability score and INT show significant increase in probability score. * With curved bracket indicates significant Tracking x Treatment interaction. * With straight bracket indicates significant effect of Treatment. ***E***, In ST, the probability to press is significantly reduced while poking is increased. In INT, both pressing and poking is significantly reduced. * Indicates significant effect of Treatment. ***F***, Latency to collect the pellet after each trial is not changed in ST but is significantly increased in INT. ***G***, Total behavior (presses + pokes) was unaffected by rimonabant in ST but decreased in INT rats. ***H***, In ST, the number of lever presses is significantly reduced while poking is increased. In INT, both pressing and poking is significantly reduced.

**Figure 4:**
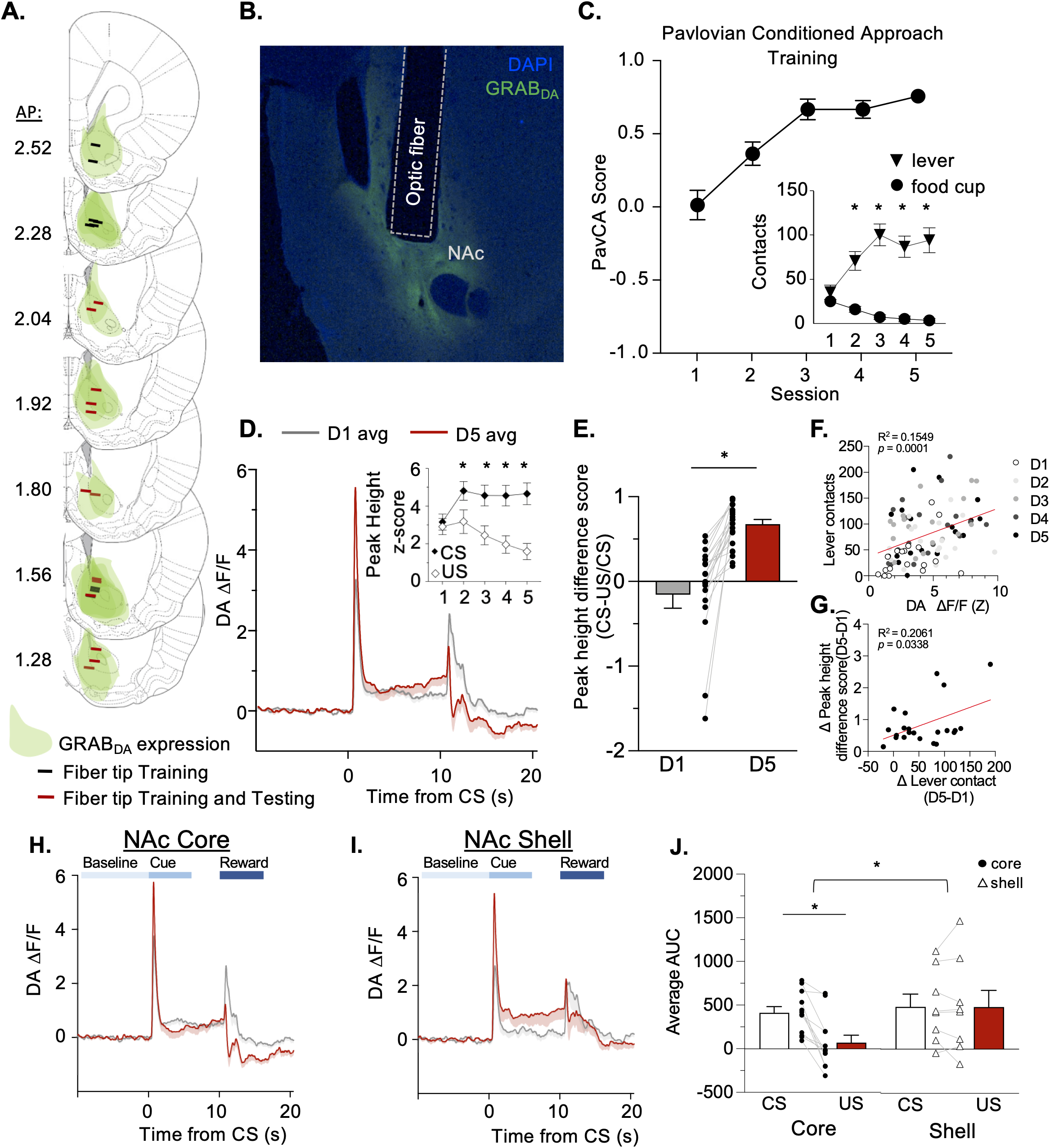
CS-evoked dopamine increases with sign-tracking behavior across Pavlovian training. ***A***, Placements of fiber optics where the GRAB_DA_ signal was recorded. Black and red fiber placements were used in the analysis of training data whereas only red fibers were used in analysis of Test data. ***B***, Example image showing optic fiber placement in the NAc and GRAB_DA_ expression in NAc core. ***C***, PavCA mean ± SEM of sign-trackers who show significantly higher presses than pokes throughout training (inset). * Indicates significant within-subjects repeated measures t-test comparing presses vs. pokes. ***D***, Mean GRAB_DA_ signal of 22 rats on D1 vs D5 of PLA. ***D inset,*** the peak height of CS-evoked dopamine is significantly higher than US dopamine on days 2-5 of training. Data from 18 rats, see methods * Significant within-subjects repeated measures t-test comparing CS vs US peak heights. **E,** We calculated the Peak height Difference Score ((CS peak height - US peak height) / CS peak height)) for each animal. A positive score indicates the CS peak is larger than the US peak. There was a significant change in this relationship over training. * Indicates main effect of Session. ***F,*** Session-by-session correlation between number of lever contacts and CS-evoked peak height for each of the five days of training. Each point represents an individual rat’s number of lever contacts and average CS-evoked peak height per session. ***G,*** Correlation between the change in NAc DA peak height difference score (D5 minus D1) and the change in lever contacts (D5 minus D1) for each rat. ***H,*** Mean GRAB_DA_ signal of n= 12 rats recorded from the NAc core of D1 (gray) vs. D5 (red) of training. ***I,*** Mean GRAB_DA_ signal of n=10 rats recorded from the NAc shell of D1 (gray) vs. D5 (red) of training. ***J,*** Area under the curve (average of D1 and D5) analysis showing differences in region (core vs. shell) and time epoch (CS vs. US) * With curved bracket indicates significant Subregion x Epoch interaction. * With straight bracket indicates significant effect of Epoch.

**Table 2:**

| Effect | Degrees Of Freedom | Lever |  |  |  |  |  | Foodcup |  |  |  |  |  |
| --- | --- | --- | --- | --- | --- | --- | --- | --- | --- | --- | --- | --- | --- |
|  |  | Contact |  | Latency |  | Probability |  | Contact |  | Latency |  | Probability |  |
|  |  | F | *p* | F | *p* | F | *p* | F | *p* | F | *p* | F | *p* |
| Session | (4,68) | 9.649 | <0.001 | 22.185 | < 0.001 | 18.602 | < 0.001 | 7.644 | < 0.001 | 3.610 | 0.01 | 4.146 | 0.005 |
| Tracking | (1,17) | 0.763 | 0.395 | 0.475 | 0.500 | 2.052 | 0.170 | 17.49 | <0.001 | 27.433 | <0.001 | 25.23 | <0.001 |
| Session x Tracking | (4,68) | 1.645 | 0.173 | 2.681 | 0.039 | 3.673 | 0.009 | 9.811 | < 0.001 | 9.049 | < 0.001 | 11.458 | < 0.001 |

Prior to the 6^th^ and 7^th^ reinforced PLA sessions, we gave rats counterbalanced intra-VTA infusions of rimonabant or vehicle. Based on previous findings (Bacharach, 2018), we predicted that intra-VTA rimonabant would reduce measures of sign-tracking behavior. Indeed, we found that blocking VTA CB1R decreased the PavCA score selectively for ST, but not INT rats (**Fig. 3C**). We found a main effect of Tracking (F(1,17) = 16.94, *p* = 0.001) and a Treatment x Tracking interaction (F(1,17) = 10.67, *p* = 0.005). This interaction was driven by a significant decrease in PavCA in the sign-trackers (*t*_(10)_ = 3.272, *p* = 0.008, Cohen’s d = 0.78) and a non-significant increase in PavCA in INT rats (*t*_(7)_ = -1.544, *p* = 0.167, Cohen’s d = 0.43).

Out of the three scores that comprise the PavCA index, the probability score was most affected by intra-VTA rimonabant injections (**Fig. 3D**). We found a main effect of Tracking (F(1,17) = 19.63, *p* < 0.001) and a Treatment x Tracking interaction (F(1,17) = 24.40, *p* < 0.001). Compared to vehicle, ST rats showed a significant decrease (*t*_(10)_ = 4.509, *p* = 0.001, Cohen’s d = 0.76) whereas INT showed a significant increase (*t*_(7)_ = -2.646, *p* = 0.033, Cohen’s d = 0.61) in the probability score when VTA CB1R signaling was decreased. To further understand the change in probability score, we compared the probability to poke vs. the probability to press across treatment conditions within each tracking group (**Fig. 3E**). Sign-trackers showed a main effect of Response (F(1,10) = 48.31, *p* < 0.001) and a Treatment x Response interaction (F(1,10) = 20.332 *p* = 0.001), suggesting that rimonabant differentially affected pressing vs. poking behavior. Sign-trackers showed a significant reduction in the probability to lever press (*t*_(10)_ = 4.734, *p* < 0.001, Cohen’s d = 0.58) and an increase in the probability to poke, though the latter wasn’t significant (*t*_(10)_ = -1.573, *p* = 0.147, Cohen’s d = 0.44). Intermediates showed a main effect of Treatment (F(1,7) = 15.16, *p* = 0.006) and Treatment x Response interaction (F(1,7) = 7.00, *p* = 0.033). Intermediates showed a reduction in the probability to press (*t*_(7)_ = 2.950, *p* = 0.021, Cohen’s d = 0.91) and a strong reduction in the probability to poke (*t*_(7)_ = 4.194, *p* = 0.004, Cohen’s d = 1.49). At first pass, these data suggest that CB1R blockade reduces all behavior in intermediate rats, this treatment instead shifts sign-tracking rats away from lever directed and towards food cup directed behaviors.

As a measure of general task engagement and motivation to consume a reward we measured the latency to collect the pellet once it was delivered for each trial (**Fig. 3F**). We found a main effect of Treatment (F(1,17) = 12.65, *p* = 0.002) and a Treatment x Tracking interaction (F(1,17) = 8.73, *p* = 0.009). Rimonabant had no effect on the latency to collect the pellet in ST rats (*t*_(10)_ = -0.686, *p* = 0.51, Cohen’s d = 0.13), suggesting that the changes in ST rat’s lever and food cup approach reported above were not due to decreased task engagement. In contrast, intra-VTA rimonabant increased the latency to collect the pellet in INT rats (*t*_(7)_ = -3.212, *p* = 0.015, Cohen’s d = 1.21), suggesting that their motivation to consume the pellets may have In part contributed to overall reduced lever and food cup approach during the cue. Regardless of latency to collect the pellet, all rats tested still ate 100% of the pellets delivered during the task, confirming there were no deficits in consummatory behavior arising from VTA CB1R manipulation.

To further examine this possibility, we next determined whether rimonabant decreased overall task engagement or motivation to consume food reward. We examined total behavior (lever contacts + foodcup contacts) during testing and found a main effect of Treatment (F(1,17) = 19.16, *p* < 0.001) and a Treatment x Tracking interaction (F(1,17) = 6.82, *p* = 0.018) (**Fig. 3G**). This interaction was driven by a significant decrease in contacts in INT rats (*t*_(7)_ = 4.741, *p* = 0.002, Cohen’s d = 1.22). These data indicate that intra-VTA rimonabant injections did not affect overall levels of conditioned responding in ST rats, but this treatment blunted behavior overall in INT rats. We further probed this interaction by investigating lever vs. foodcup contacts in each tracking group (**Fig. 3H**). Similar to the probability data, ST rats showed a main effect of Response (F(1,10) = 24.732, *p* = 0.001) and a Treatment x Response interaction (F(1,10) = 10.661, *p* = 0.0009). This interaction was driven by a decrease in lever contacts (*t*_(10)_ = 3.167, *p* = 0.01, Cohen’s d = 0.50) and an increase in foodcup pokes (*t*_(10)_ = -1.273, *p* = 0.116, Cohen’s d = 0.44). In contrast, INT rats showed a reduction in both responses; main effect of Treatment (F(1,7) = 22.473, *p* < 0.002). In summary, ST had similar amounts of total approach behavior in which a reduction in lever-directed behavior was compensated for by an increase in foodcup-directed behavior. INT rats still engaged in both types of responding, but both responses were decreased.

As a control, we determined whether locomotor activity changed with intra-VTA rimonabant injections. We gave an open-field test to a subset of ST and INT rats and found no differences in locomotor activity when rats were given intra-VTA vehicle or rimonabant infusions (distance travelled in meters: Veh mean ± SEM = 19.65, 1.44; Rimo mean ± SEM = 18.48, 0.86) . There were no main effects of Tracking, Treatment, nor a Tracking x Treatment interaction (Max F = 1.147, Min *p* = 0.307).

Taken together, CB1R inhibition in the VTA causes divergent behavioral response profiles across ST and INT rats. Rimonabant in the VTA caused a decrease in all appetitive motivated behaviors measured in the INT rats. In contrast, rimonabant only decreased behavior directed towards the lever cue in ST rats, while increasing the amount of cue-induced food cup behaviors. Thus, CB1R receptor blockade decreases sign-tracking behavior leading to more balanced levels of lever- and foodcup-directed behaviors.

### CS-evoked dopamine increases with sign-tracking behavior across Pavlovian training

In order to understand rimonabant’s effect on cue-evoked dopamine in the sign-trackers, we measured GRAB_DA_ signals in the NAc core and shell while manipulating CB1R in the VTA. We infused the fluorescent dopamine sensor, GRAB_DA_, and implanted fiber optic cannula targeting the NAc in separate group of 22 sign-tracking rats (**Fig. 4A, B**). Rats acquired a sign-tracking response as previously observed (PavCA: main effect of Session (F(4,84) = 33.630, *p* < 0.001) (**Fig. 4C**). Further, when examining pressing and poking contacts, ST rats showed main of effects of Session (F(4,84) = 5.229, *p* < 0.001) and Response (F(1,84) = 48.689, *p* = 0.003), and a Session x Response interaction (F(4,84) = 19.172, *p* < 0.001) where pressing behavior was significantly greater than poking behavior on D2-D5 of training (*p’s* < .0001) (**Fig. 4C inset**).

Voltammetry studies establish that ST, but not GT, show a transfer of DA transients from the US to CS over Pavlovian training (Clark et al., 2013; Flagel et al., 2011; Saddoris et al., 2016) which is interpreted as signal supporting enhanced motivational value of the CS in ST rats. We examined the NAc GRAB_DA_ signal across training (**Fig. 4D, inset**) and found main effects of Session (F(4,68) = 5.044, *p* = 0.001) and Epoch (CS,US; F(1,17) = 36.414, *p* < 0.001) and a Session x Epoch interaction (F(4,68) = 19.022, *p* < 0.001). CS evoked NAc GRAB_DA_ signal was greater than US evoked GRAB_DA_ signal on D2-D5 of training (*p’s* < 0.0007). This data reflect the classic signature of a dopamine response to transfer from the US to the CS across conditioning (Day et al., 2007; Schultz et al., 1997).

We next compared the relationship of CS- and US-evoked on GRAB_DA_ signals for each rat across D1 and D5. We calculated the peak height difference score (peak height: CS-US/CS) on D1 versus D5 (**Fig. 4E**). We found that there was a significant increase in this score on D5 compared to D1 of training (*t*_(21)_ = -5.771, *p* < 0.001, Cohen’s d = 1.53), demonstrating that on GRAB_DA_ signals to the US were transferred to the CS in ST over the course of training.

On a session-by-session basis we observe the magnitude of sign-tracking (lever contact) positively correlates with CS-evoked NAc DA peak heights across training (sessions 1-5, n is rat average per session; R^2=^0.155, *p* = 0.0001, **Fig. 4F**). Furthermore, the increase in sign-tracking across training (D5 minus D1 lever contacts) correlates positively with the change in peak height difference scores (CS-US/CS; D1-D5) (R^2^ = 0.2061, *p* = 0.0338) (**Fig. 4G**). Together these data suggest that greater escalation of sign-tracking is associated with greater CS relative to US DA signals across conditioning.

Due to known differences in dopamine release and dynamics in NAc core vs. shell (Cacciapaglia et al., 2012; Saddoris et al., 2013; West & Carelli, 2016; Zahm, 1999), we examine region-specific NAc GRAB_DA_ signals across conditioning (**Fig 4H, I**). We examined the area under the curve between two time epochs (Cue (CS): 5s post lever insertion, Reward (US): 5s post lever retraction), which reveals main effect of Epoch (CS > US, F(1,18) = 12.97, *p* = 0.002) and an Epoch x Region interaction (F(1,18) = 12.76, *p* = 0.002, **Fig. 4J**), which is driven by greater NAc core CS vs US responses (*t*_(11)_ = 5.220, *p* < 0.001, Cohen’s d = 1.26) and greater US GRAB_DA_ responses in NAc shell than NAc core (*t*_(18)_ = 2.184, *p* = 0.042, Cohen’s d = 0.94). These results confirm that cue-evoked NAc GRAB_DA_ signal increases to and correlated lever directed behavior in sign-tracking rats, consistent with prior reports (Day et al., 2007; Flagel et al., 2011; Clark et al., 2013; Saddoris et al., 2016). We also provide evidence that dopamine is released differentially in the core versus shell during training.

### Intra-VTA rimonabant primarily affects dopamine encoding of outcomes rather than cues

We used thirteen ST rats in our analysis of test data. Nine rats were excluded from test analysis due to off target cannula placement in the VTA infusion site (n=4), excessive damage at VTA infusion site (n=2), or a loss of signal during testing (n=3). Similar to our prior behavioral findings, we found intra-VTA rimonabant significantly decreased the PavCA score of ST rats (*t*_(12)_ = 3.476, *p* = 0.005, Cohen’s d =0.82) (**Fig. 5A**). The decrease in PavCA was accompanied by similar reductions in the preference, latency, and probability scores (vehicle vs. rimonabant; *p’s =* 0.019, 0.005, 0.002, respectively).

**Figure 5:**
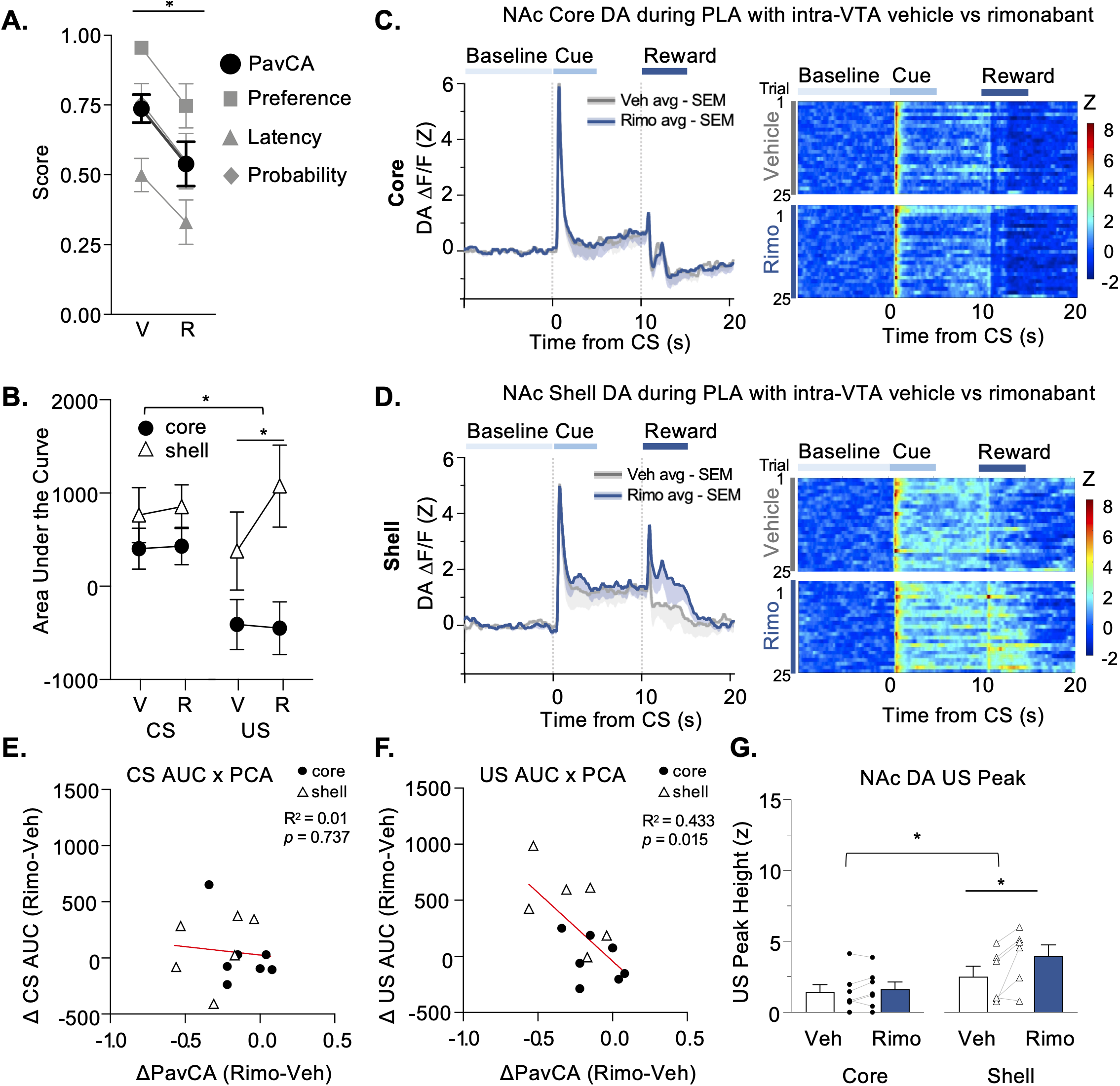
Intra-VTA rimonabant primarily affects dopamine encoding of outcomes rather than cues. ***A***, Rimonabant significantly reduces PavCA score in ST rats, as well as preference, latency, and probability scores. * Indicates main effect of Session. ***B***, Area under the curve for the 5s CS period vs. 5s US period in animals recorded in NAc core vs. shell under vehicle or rimonabant conditions. * With curved bracket indicates significant Subregion x Treatment interaction. * With straight bracket indicates significant effect of Treatment. ***C left,*** Mean -SEM shading of GRAB_DA_ signal in the NAc core in vehicle (gray) vs. rimonabant (blue) conditions. ***C right*,** trial by trial heatmap of average GRAB_DA_ signal in NAc core over vehicle and rimonabant conditions. ***D left,*** Mean -SEM shading of GRAB_DA_ signal in the NAc shell in vehicle (gray) vs. rimonabant (blue) conditions ***D right*,** trial by trial heatmap of average GRAB_DA_ signal in NAc shell over vehicle and rimonabant conditions. ***E,*** Correlation between the change in area under the curve for CS epoch (Rimo minus vehicle) with change in the PavCA score (rimo minus vehicle). ***F,*** Correlation of the change in area under the curve for US epoch (rimo minus vehicle) with change in the PavCA score (rimo minus vehicle). ***G,*** US peak height analysis showing differences in region (core vs. shell) and treatment (core vs. shell) * With curved bracket indicates significant Subregion x Treatment interaction. * With straight bracket indicates significant effect of Treatment.

Due to known differences in dopamine release and dynamics in NAc core vs. shell in training and consistent with prior reports (Cacciapaglia et al., 2012; Saddoris et al., 2013; West & Carelli, 2016; Zahm, 1999), we include NAc Subregion as a statistical factor in addition to Epoch and Treatment (Rimonabant, Vehicle). We first examined the AUC during the two epochs and found a significant NAc Subregion x Epoch x Treatment interaction (F(1,11) = 5.311, *p* = 0.042) (**Fig. 5B**). To follow up, we analyzed the total GRAB_DA_ signal during the US period and found a main effect of Treatment (F(1,11) = 7.919, *p* = 0.017), and a Treatment x Subregion interaction (F(1,11) = 9.996, *p* = 0.009). This interaction was driven by increases in GRAB_DA_ signals specifically in the NAc shell during the US reward period (*t*_(5)_ = -3.257, *p* = 0.023, Cohen’s d = 0.66). We present NAc subregion specific traces and heatmaps in **Fig. 5C** and **5D**.

We next probed if the rimonabant-induced NAc DA signal changes were associated with behavioral changes in Pavlovian lever autoshaping. We correlated changes in dopamine signals during the CS and US periods (AUC^Rimo^ - AUC^Veh^) with changes in PavCA score (PavCA^Rimo^ -PavCA^Veh^). We found that treatment-induced changes in NAc GRAB_DA_ signals during the CS period did not correlate with changes in PavCA score (R^2^ = 0.01, *p* = 0.734) (**Fig. 5E**). However, treatment-induced changes in NAc GRAB_DA_ signals during the US period were negatively correlated with changes in the PavCA score (R^2^ = 0.433, *p* = 0.015) (**Fig. 5F**). This suggests that the more intra-VTA rimonabant shifted behavior towards goal-tracking PavCAs (more negative scores), the greater the rimonabant-induced NAc DA signaling during the US period relative to vehicle. We found similar negative correlations between rimonabant-induced changes in US-AUC and the Preference, Latency, and Probability scores (*p’s* = 0.006, 0.100, 0.059, respectively).

Consistent with this, US peak heights were increased with rimonabant treatment (main effect of Treatment: (F(1,11) = 10.228, *p* = 0.008) and varied by NAc subregion (Treatment x Subregion interaction (F(1,11) = 5.77, *p* = 0.035) **Fig. 5G**). This was driven by increases in US peak height in the NAc shell (*t*_(5)_ = -2.735, *p* = 0.041, Cohen’s d = 0.78). However, treatment-induced changes in US peak height did not predict variability in PavCA or other behavioral measures, suggesting that the sustained US NAc DA signaling that is captured by the AUC analysis (above) relates more to behavior than the US peak.

We next examined dopamine dynamics surround the CS. On a population level, the CS peak height was not affected by intra-VTA rimonabant treatment and we observed a large amount of variability in CS peaks across NAc (**Fig. 6A**). Thus, we examined treatment-related changes in CS-evoked peak (CSpeak^Rimo^ - CSpeak^Veh^) as a function of changes in PavCA score (PavCA^Rimo^ -PavCA^Veh^), which were positively correlated (**Fig. 6B**). This suggests that the more intra-VTA rimonabant shifted behavior towards goal-tracking PavCAs (more negative PavCA difference score), the smaller the rimonabant-induced NAc DA CS peak relative to vehicle. Notably, rats that showed less of a behavioral shift (PavCA difference scores near zero) tended to have greater CS- evoked DA peaks with intra-VTA rimonabant relative to vehicle. This suggests the more rigid the behavior was between treatment sessions, the greater the CS-evoked NAc dopamine signaling when VTA CB1R signaling was disrupted. Placements of VTA infusions and NAc recording sites are given in **Fig. 6C,D.** Taken together, with inhibition of CB1R signaling in the VTA we find that shifts in behavior away from sign-tracking and towards goal-tracking (negative PavCA shifts) are associated with smaller NAc DA CS peaks (**Fig. 6B**) and greater NAc DA US processing (**Fig. 5F**).

**Figure 6.**
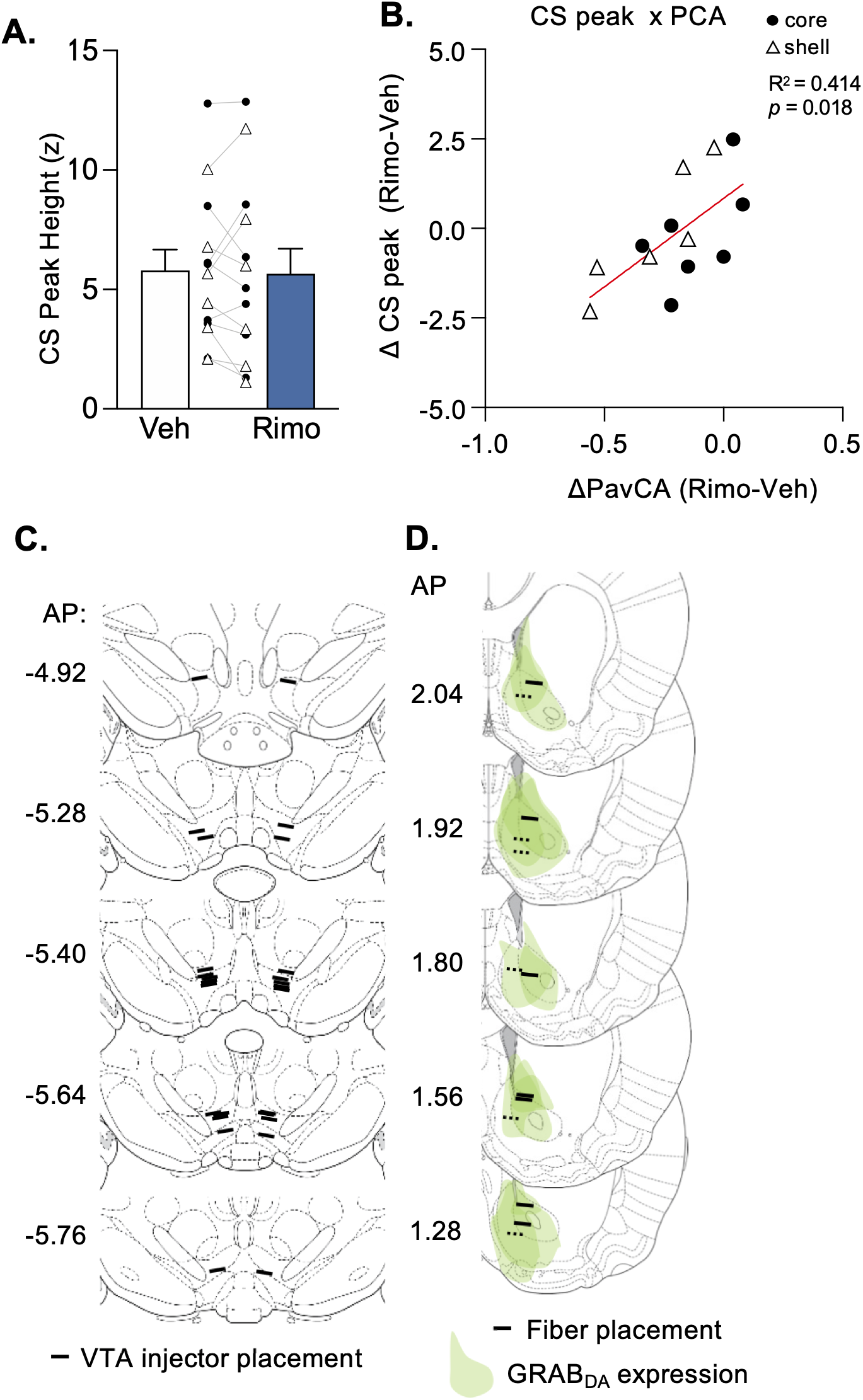
CB1R inhibition effect on CS-evoked peak dynamics predict magnitude of treatment effect on behavior. ***A,*** Rimonabant does not change CS-induced peak height in core or shell. ***B,*** Correlation of the change in CS-evoked peak height (rimo minus vehicle) with the change in PavCA score (rimo minus vehicle). ***C***, Site of injector tip placement in the VTA for rimonabant infusion. ***D,*** GRAB_DA_ expression and photometry recording location in NAc. Solid lines indicate recording location for NAc Core and dotted lines indicate location of NAc Shell fibers.

### CS-evoked dopamine is specific to rewarded lever

Lastly, as a control in separate groups of rats, we confirmed that the GRAB_DA_ signals in the NAc are specific to the learned association between the CS and the US. We trained a separate group of fiber implanted, NAc GRAB_DA_ expressing rats (**Fig. 7A**) in a 2-lever Pavlovian autoshaping procedure. Here, a CS+ lever predicted food reward and a separate CS- lever predicted no reward. We examined total behavior (lever and food cup contact) during the CS+ versus CS- periods across training (**Fig. 7B**) and found a main effect of Cue (F(1,5) = 11.939, *p* = 0.018) and a Cue x Session interaction (F(4,20) = 11.325, *p* < 0.001). Rats showed more approach during the CS+, compared to the CS-presentations on D5 of training (*t*_(5)_ = 3.598, *p* = 0.016, Cohen’s d = 1.88). This confirms that by D5 of training, the rats could behaviorally discriminate between a reinforced from nonreinforced cue. On D5 of training, we found the CS evoked peak height was greater for the CS+ lever than the CS (*t*_(5)_ = 2.947, *p* = 0.032, Cohen’s d = 0.79; **Fig. 7 C,D**). We also trained a sperate group of fiber implanted, NAc GRAB_DA_ expressing rats (n=4) rats in an unpaired control condition in which we delivered the same number of food pellets, but lever extension was delivered pseudorandomly during the ITI period and was never explicitly paired with pellet delivery. Unpaired rats displayed low levels of lever and foodcup approach during lever extension periods throughout training **(Fig. 7E**). Rats did not develop conditioned responses to either the lever or foodcup (D1 vs. D5 (*t*_(3)_ = 1.023, *p* = 0.382) (**Fig. 7E inset left**). Despite not showing conditioned lever or foodcup approach, rats were still engaged in the task and collected the pellet faster on D5 than D1 of training (*t*_(3)_ = 3.696, *p* = 0.034, Cohen’s d = 2.56) (**Fig. 7E inset right**). In this unpaired condition, we observed very low levels of lever-associated NAc GRAB_DA_ signals which diminished, though not significantly, as training progressed (D1 vs. D5 *t*_(3)_ = 2.425, *p* = 0.119) (**Fig. 7F**). In the unpaired rats, NAc GRAB_DA_ signals to the CS on D1 likely represent a salience or novelty signal (Horvitz et al., 1997; Mirenowicz & Schultz, 1994), which goes away over time as rats become familiar with the lever extension and retraction. There was reliable NAc GRAB_DA_ signal to the US food delivery (**Fig. 7G**), that remained unchanged from D1 to D5 of training (*t*_(3)_ = -1.594, *p* = 0.209) (**Fig. 7G inset**), confirming the unpredictable delivery of the US in unpaired rats. Our results indicate that the NAc GRAB_DA_ signal observed to the CS and US during the paired, single-lever condition arise due to rats learning and maintaining the CS-US relationship.

**Figure 7.**
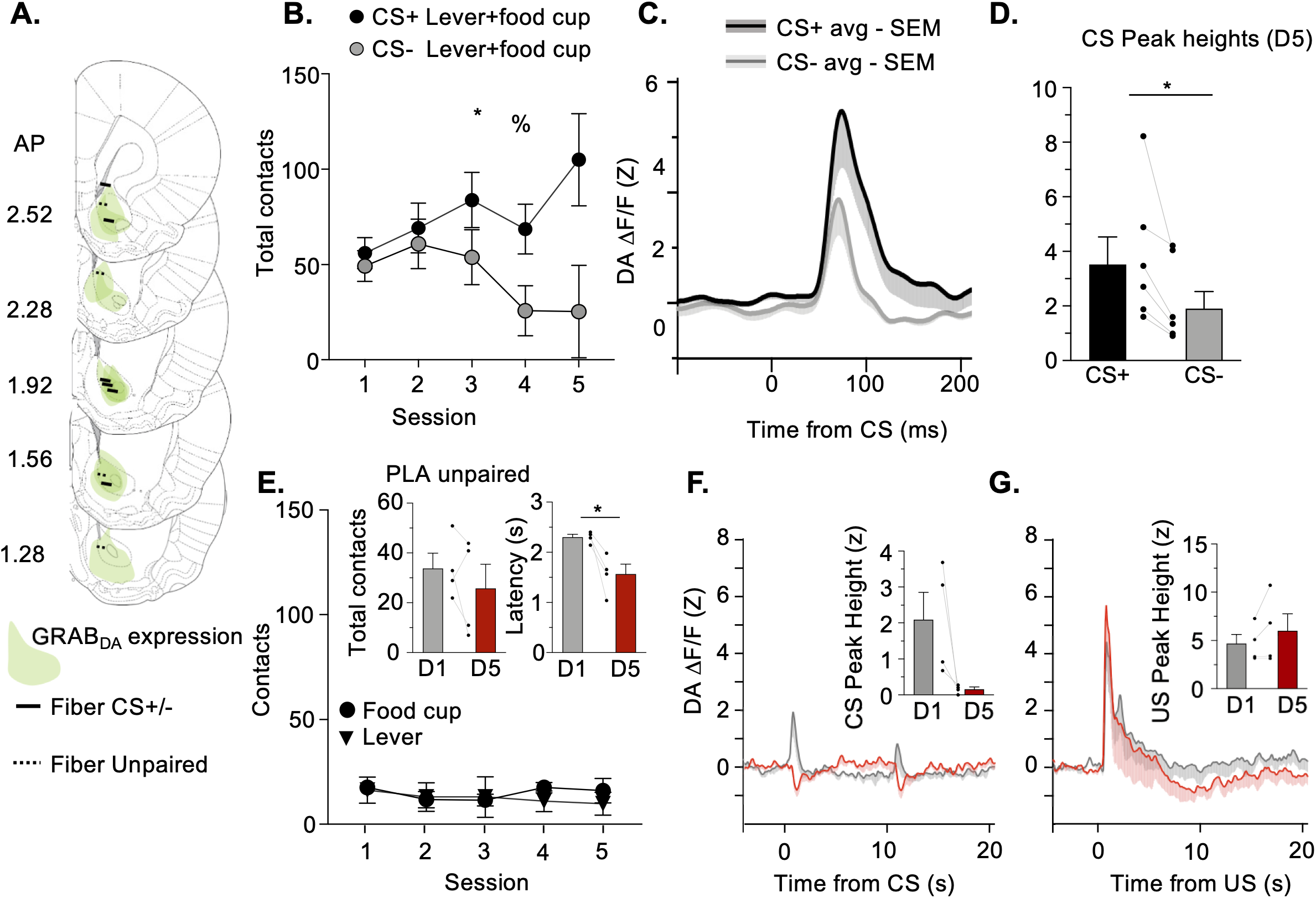
CS-evoked dopamine is specific to rewarded lever. ***A***, Histological placements of photometry fibers in the NAc. Two separate groups of rats were used for training in 2-lever PLA or Unpaired PLA. 6 rats were run in the 2-lever program and their fiber placements are indicated by the solid black line. Four rats were run in the unpaired program, placements indicated with dashed line. ***B***, Rats show behavioral discrimination (presses + pokes) between the CS+ and CS- lever. * Main effect of Session; % Significant Session x Response interaction. ***C****, **D*** The CS-evoked GRAB_DA_ signal is significantly higher to the rewarded than the unrewarded lever by day 5 of training. * Indicates main effect of Stimulus. ***E,*** Rats in the Unpaired condition display low amounts of behavior towards both the lever or foodcup. *Left inset*: total behavior does not increase from D1 to D5 suggesting no conditioned behavior has emerged. *Right inset*: Latency to collect the pellet reduces over training, suggesting rats are still engaged and food-motivated during the task. * Indicates main effect of Session. ***F***, GRAB_DA_ responses to the lever on D1 vs D5 of training: A CS-evoked excitatory response to the lever is not established. ***G***, GRAB_DA_ signal to pellet delivery remains high across training.

## DISCUSSION

Here, we demonstrate that optogenetic inhibition of VTA-NAc core dopamine projections reduces lever approach, specifically in sign-tracking rats. While intermediate rats also display lever approach, VTA-NAc core DA terminal inhibition did not affect this behavior, nor did it affect the preferred food cup response in goal-tracking rats. Hence, VTA→NAc core dopamine release during the cue period is necessary for the enhanced cue attraction specifically seen in sign-tracking rats. Having confirmed the importance of VTA-NAc core DA projections in ST rats, we next examined whether disrupting VTA CB1R signaling similarly decreased lever approach in sign-tracking rats. We compared the effects of intra-VTA rimonabant injections in ST and intermediate rats because both display lever directed approach but differently engage VTA-NAc core. We found that intra-VTA infusions of the CB1R inverse agonist, rimonabant, resulted in different response profiles between ST and INT rats. In INT rats, decreasing CB1R signaling decreased all appetitive motivated behaviors that we measured. Both lever and food cup responses decreased and the latency to collect the reward increased. In ST rats, intra-VTA rimonabant selectively reduced lever approach while increasing food-cup approach, shifting the rats away from sign-tracking (lever directed) and towards goal-tracking (food-cup behaviors). These results suggest that intact VTA CB1R signaling biases behavior towards the lever and away from the foodcup in ST rats. We predicted that the VTA rimonabant-induced decrease in cue attraction would be associated with disrupted cue-triggered dopamine signals downstream in the NAc. While we did not observe overall effects of VTA CB1R inhibition on CS evoked NAc DA signaling, VTA rimonabant-induced behavioral shifts away from sign-tracking were positively correlated with changes to cue-evoked NAc core GRAB_DA_ signals. Rats that showed the greatest changes in behavior with intra-VTA rimonabant (away from ST) tended to have decreased NAc DA CS peaks during rimonabant relative to vehicle sessions. In contrast, rats that showed less of a behavioral shift (i.e., more rigid behavior between sessions) tended to have greater CS-evoked DA peaks during rimonabant relative to vehicle sessions.

We also found that intra-VTA rimonabant increased NAc GRAB_DA_ signals during the reward period and that this effect was driven by increased US DA signaling in the NAc shell. VTA rimonabant-induced behavioral shifts away from sign-tracking were negatively correlated with changes to US-evoked NAc core GRAB_DA_ signals. That is, rats that showed the greatest changes in behavior (away from ST) with intra-VTA rimonabant tended to have increased NAc DA US peaks during rimonabant relative to vehicle sessions. In contrast, rats that showed less of a behavioral shift with intra-VTA rimonabant tended to have lower US-evoked DA peaks during rimonabant relative to vehicle sessions. Our results demonstrate decreasing VTA CB1R signaling decreases sign-tracking and increases goal-tracking approach, potentially by suppressing cue-evoked, and enhancing reward-evoked NAc dopamine signals.

Prior pharmacology and voltammetry tracking studies have established NAc core DA as necessary for sign-tracking (Flagel et al., 2011; Saunders & Robinson, 2012; Clark et al., 2013; Fraser & Janak, 2017). Consistent with manipulations of NAc dopamine receptors, we show that optogenetic inhibition of VTA-NAc core dopamine projections during lever presentation similarly reduces lever approach in ST rats, but not in intermediate rats that also display lever approach. Our results using Pavlovian conditioning are consistent with a substantial instrumental literature showing that CB1R signaling maintains both cued food-seeking and drug-seeking behaviors (McLaughlin et al., 2003; De Vries & Schoffelmeer, 2005; Ward & Dykstra, 2005; Economidou et al., 2006; Salamone et al., 2007; Ward et al., 2007; Justinova et al., 2008; de Bruin et al., 2011; Oleson et al., 2012; Schindler et al., 2016). Consistently, sign-trackers and intermediates show decreases in lever approach when we disrupt VTA CB1R signaling. However, in intermediate rats this attenuating effect was not specific to the lever, as food cup responding and task engagement were generally reduced, consistent with prior rimonabant manipulations (McLaughlin et al., 2003; Ward & Dykstra, 2005; McLaughlin et al., 2006; Salamone et al., 2007; Ward et al., 2007; Oleson et al., 2012). However, this was not the case in sign-trackers that decreased lever approach but showed no differences in overall task engagement and tended to increase food cup responding when VTA CB1R signaling was disrupted. Thus, blocking VTA CB1R signaling rebalanced lever- and foodcup-directed behavior, such that a decrease in lever approach was compensated for by an increase food cup approach to result in a significant decrease in the PavCA score. The eCB system is also involved in maintaining Pavlovian cue-reward associations. Systemic blockade of CB1R signaling disrupts cue-driven approach behaviors in Pavlovian tasks (Bacharach et al., 2018; Gheidi et al., 2020). The present findings using region specific manipulation of CB1R provide evidence that the VTA is a site of endocannabinoid action to promote sign-tracking and confirm our prior work showing systemic rimonabant dose-dependently reduces sign-tracking and the attracting and reinforcing properties of the lever cue (Bacharach et al., 2018).

We found evidence that manipulating VTA CB1R signaling influences dopamine release during the US/reward period that varies by NAc subregion. There are anatomical and functional differences between the core and shell (Jones et al., 1996; Zahm, 1999). In cued-instrumental and Pavlovian settings, NAc core dopamine release is associated with the predictive value of a cue (Roitman et al., 2004; Cacciapaglia et al., 2012; Hart et al., 2014; Saddoris et al., 2015; Stelly et al., 2021). In addition to the predictive value of this dopamine signal, core dopamine also carries incentive motivational value (Berridge, 1996; Flagel et al., 2011; Saunders & Robinson, 2012; Saunders et al., 2013). Our photometry and optogenetic data from the NAc core corroborate these findings in that NAc core is necessary for cue-approach in sign-trackers and we see strong cue-induced dopamine under both vehicle on rimonabant conditions. We find both NAc core DA inhibition and VTA CB1R antagonism decrease sign-tracking. In our photometry recordings, the relationship between VTA CB1R signaling and cue-evoked DA signaling spans recordings from both the NAc core and shell. We observe consistent effects of VTA CB1R antagonism on PavCA scores across our pharmacology experiment (**Fig. 3**) when rats are untethered, and our combined pharmacology and photometry experiment (**Fig. 5**) when rats are tethered. This suggests this technical difference of tethering rats, which does moderately slow responding, did not interfere with VTA treatment effects on behavior. We do not observe NAc region-specific differences in effects of VTA CB1R antagonism on cue-evoked DA signaling. Overall VTA CB1R antagonism does not affect CS-evoked NAc DA, but variability in the effects of rimonabant on behavior do relate to magnitude of treatment effects on CS evoked NAc DA. This relationship suggests VTA CB1R signaling may influence NAc DA encoding of predictive or incentive motivational properties of the CS and thus bias behavior towards cues.

The NAc shell DA is more heavily involved in representing contextual elements associated with reward seeking, tracking reinforcer value, and representing reward-guided motivation (Bossert et al., 2007, 2012; Cacciapaglia et al., 2012; Cruz et al., 2014; West & Carelli, 2016; Valyear et al., 2020). Additionally, NAc shell dopamine release is less temporally restricted to cue onset, and differential dopamine dynamics are observed during cue-onset, the entire cue period, and the US period (Saddoris et al., 2015, 2016). We find that VTA CB1R blockade *increases* NAc shell dopamine signaling during the US period and is associated with increased food cup exploration, which may give rise to changes in US processing and/or reinforcer value representation.

The present study used almost exclusively female rats, and thus was not designed to test for sex differences. Here we find that greater shifts in sign-tracking are associated with decreased NAc CS DA during intra-VTA rimonabant vs vehicle conditions. Yet the effect of rimonabant was more subtle than in our prior study using voltammetry in male rats (Oleson, Beckert et al. 2012), as group effects of VTA rimonabant in the present study were not evident. Instead, the decrease in CS-evoked NAc DA was greatest in female rats that shifted away from sign-tracking the most. It is likely differences in the sex of rats and DA recording techniques are the most likely factors influencing different observations between studies. We used female rats for several reasons. First, we and others see evidence for increased propensity for females to sign-track (Pitchers et al., 2015; Kochli et al., 2020; Madayag et al., 2017). Second, in our systemic rimonabant study, we observed greater CB1R manipulation effect sizes in females compared to males on the attracting and reinforcing properties of cues (Bacharach et al., 2018). Consistent with these behavioral and pharmacological findings, a recent study shows higher conditioned responding in females compared to males (Lefner et al., 2022). Females in that study showed lower US-evoked NAc DA responses compared to males during Pavlovian conditioning. This latter result has relevance for the present findings, which point towards a role for VTA CB1R regulation of behavioral and dopaminergic cue bias observed in female rats. Future studies designed to probe sex differences would be needed to determine if the present findings are sex-specific or not.

Here we examine effects of VTA CB1R signaling on NAc dopamine during Pavlovian behavior, while prior work examining such effects used instrumental tasks. Importantly, prior instrumental studies used short, fixed inter trial intervals (10-20 s), which has implications for CB1R receptor involvement in dopamine neuron activity. Dopamine is intimately involved in interval timing (Buhusi & Meck, 2005; Mikhael & Gershman, 2019). Oleson et al. (2012) found that there was larger cue-induced dopamine release in a fixed interval, which affected more by rimonabant, versus a variable interval task. Here we used a Pavlovian lever autoshaping task with a much longer 90s variable intertrial interval. This long and unpredictable interval between trials may limit our ability to see group differences in CS-evoked NAc DA changes with CB1R manipulations. Regardless, the present study revealed the greatest rimonabant-induced decrease in CS-evoked NAc DA were associated with the largest decreases in Pavlovian sign-tracking. This finding corresponds with our prior work indicating VTA CB1R signaling gates cue-evoked NAc DA to regulate instrumental motivational processes (Oleson, Beckert et al. 2012).

Together, our results suggest that both VTA CB1R signaling and NAc core DA support the maximal expression of sign-tracking behavior. Disrupting VTA CB1R signaling rebalances behavior away from sign-tracking and towards goal-tracking. Further, the intra-VTA rimonabant induced shift in behavior of ST rats is related to decreases in CS- and increases in US-evoked NAc shell dopamine signaling, indicating that VTA CB1R receptors are involved in maintaining NAc representations of the CS-US relationship. Our results suggest that CB1R signaling in the VTA maintains the conditioning-dependent behavioral and NAc DA bias towards cues relative to outcomes in sign-tracking rats, potentially by downregulating reward-related NAc shell DA signaling. Future causal role studies are needed to test the necessity of NAc shell DA for mediating the balance between CS and US directed behaviors in lever autoshaping.

## Conflict of Interest

The authors declare that the research was conducted in the absence of any commercial or financial relationships that could be construed as a potential conflict of interest.

## Acknowledgements

This work was supported by a McKnight Memory and Cognitive Disorders Award (McKnight Foundation; DC), and two National Institute on Drug Abuse (NIDA) grant R01DA043533 (DC), F31DA050367 (SB) and the Department of Anatomy and Neurobiology at the University of Maryland, School of Medicine. The funders had no role in the study design, data collection, and analysis, decision to publish, or preparation of the manuscript. We thank Jessie Feng for technical assistance with GRAB_DA_ sensors and the University of Maryland, Baltimore Animal Care Facility staff for colony maintenance. We thank Michael McDannald for insightful comments and discussions on this work. We thank Asaf Keller and the Department of Anatomy and Neurobiology for sharing photometry equipment.

## Data Availability Statement

The raw data supporting the conclusions of this article will be made available by the authors, without undue reservation, to any qualified researcher.

## Ethics Statement

The animal study was reviewed and approved by University of Maryland, School of Medicine Institutional Animal Care and Use Committee.

## Author Contributions

DC, SB and JC conceived, developed and supervised the project. SB acquired the data. SB, DM and CS analyzed the data. SB and DC designed the experiments, interpreted the data, and wrote the manuscript. FS and YL produced and provided the dopamine sensor, GRAB_DA_, used for fiber photometry. All authors contributed to manuscript revision and approved the submitted version.

## Notes

### Competing Interest Statement

The authors have declared no competing interest.

### Summary of Updates

This revision contains new analyses and additional text revisions requested in first round of peer review

## References

Bacharach, S. Z., Nasser, H. M., Zlebnik, N. E., Dantrassy, H. M., Kochli, D. E., Gyawali, U., Cheer, J. F., & Calu, D. J. (2018). Cannabinoid receptor-1 signaling contributions to sign-tracking and conditioned reinforcement in rats. Psychopharmacology, 235(10), 3031–3043. https://doi.org/10.1007/s00213-018-4993-6

Berridge, K. C. (1996). Food reward: Brain substrates of wanting and liking. Neuroscience & Biobehavioral Reviews, 20(1), 1–25. https://doi.org/10.1016/0149-7634(95)00033-B

Bossert, J. M., Poles, G. C., Wihbey, K. A., Koya, E., & Shaham, Y. (2007). Differential effects of blockade of dopamine D1-family receptors in nucleus accumbens core or shell on reinstatement of heroin seeking induced by contextual and discrete cues. The Journal of Neuroscience: The Official Journal of the Society for Neuroscience, 27(46), 12655–12663. https://doi.org/10.1523/JNEUROSCI.3926-07.2007

Bossert, J. M., Stern, A. L., Theberge, F. R. M., Marchant, N. J., Wang, H.-L., Morales, M., & Shaham, Y. (2012). Role of projections from ventral medial prefrontal cortex to nucleus accumbens shell in context-induced reinstatement of heroin seeking. The Journal of Neuroscience: The Official Journal of the Society for Neuroscience, 32(14), 4982–4991. https://doi.org/10.1523/JNEUROSCI.0005-12.2012

Buhusi, C. V., & Meck, W. H. (2005). What makes us tick? Functional and neural mechanisms of interval timing. Nature Reviews. Neuroscience, 6(10), 755–765. https://doi.org/10.1038/nrn1764

Cacciapaglia, F., Saddoris, M. P., Wightman, R. M., & Carelli, R. M. (2012). Differential dopamine release dynamics in the nucleus accumbens core and shell track distinct aspects of goal-directed behavior for sucrose. Neuropharmacology, 62(5–6), 2050–2056. https://doi.org/10.1016/j.neuropharm.2011.12.027

Cheer, J. F., Marsden, C. A., Kendall, D. A., & Mason, R. (2000). Lack of response suppression follows repeated ventral tegmental cannabinoid administration: An in vitro electrophysiological study. Neuroscience, 99(4), 661–667. https://doi.org/10.1016/s0306-4522(00)00241-4

Cheer, J. F., Wassum, K. M., Heien, M. L. A. V., Phillips, P. E. M., & Wightman, R. M. (2004). Cannabinoids enhance subsecond dopamine release in the nucleus accumbens of awake rats. The Journal of Neuroscience: The Official Journal of the Society for Neuroscience, 24(18), 4393–4400. https://doi.org/10.1523/JNEUROSCI.0529-04.2004

Cheer, J. F., Wassum, K. M., Sombers, L. A., Heien, M. L. A. V., Ariansen, J. L., Aragona, B. J., Phillips, P. E. M., & Wightman, R. M. (2007). Phasic dopamine release evoked by abused substances requires cannabinoid receptor activation. The Journal of Neuroscience: The Official Journal of the Society for Neuroscience, 27(4), 791–795. https://doi.org/10.1523/JNEUROSCI.4152-06.2007

Clark, J. J., Collins, A. L., Sanford, C. A., & Phillips, P. E. M. (2013). Dopamine Encoding of Pavlovian Incentive Stimuli Diminishes with Extended Training. Journal of Neuroscience, 33(8), 3526–3532. https://doi.org/10.1523/JNEUROSCI.5119-12.2013

Cruz, F. C., Babin, K. R., Leao, R. M., Goldart, E. M., Bossert, J. M., Shaham, Y., & Hope, B. T. (2014). Role of nucleus accumbens shell neuronal ensembles in context-induced reinstatement of cocaine-seeking. The Journal of Neuroscience: The Official Journal of the Society for Neuroscience, 34(22), 7437–7446. https://doi.org/10.1523/JNEUROSCI.0238-14.2014

Danna, C. L., & Elmer, G. I. (2010). Disruption of conditioned reward association by typical and atypical antipsychotics. Pharmacology Biochemistry and Behavior, 96(1), 40–47. https://doi.org/10.1016/j.pbb.2010.04.004

Day, J. J., Roitman, M. F., Wightman, R. M., & Carelli, R. M. (2007). Associative learning mediates dynamic shifts in dopamine signaling in the nucleus accumbens. Nature Neuroscience, 10(8), 1020–1028. https://doi.org/10.1038/nn1923

de Bruin, N. M. W. J., Lange, J. H. M., Kruse, C. G., Herremans, A. H., Schoffelmeer, A N. M., van Drimmelen, M., & De Vries, T. J. (2011). SLV330, a cannabinoid CB(1) receptor antagonist, attenuates ethanol and nicotine seeking and improves inhibitory response control in rats. Behavioural Brain Research, 217(2), 408–415. https://doi.org/10.1016/j.bbr.2010.11.013

De Vries, T. J., & Schoffelmeer, A. N. M. (2005). Cannabinoid CB1 receptors control conditioned drug seeking. Trends in Pharmacological Sciences, 26(8), 420–426. https://doi.org/10.1016/j.tips.2005.06.002

Economidou, D., Mattioli, L., Cifani, C., Perfumi, M., Massi, M., Cuomo, V., Trabace, L., & Ciccocioppo, R. (2006). Effect of the cannabinoid CB1 receptor antagonist SR-141716A on ethanol self-administration and ethanol-seeking behaviour in rats. Psychopharmacology, 183(4), 394–403. https://doi.org/10.1007/s00213-005-0199-9

Flagel, S. B., Clark, J. J., Robinson, T. E., Mayo, L., Czuj, A., Willuhn, I., Akers, C. A., Clinton, S. M., Phillips, P. E. M., & Akil, H. (2011). A selective role for dopamine in stimulus–reward learning. Nature, 469(7328), 53–57. https://doi.org/10.1038/nature09588

Flagel, S. B., Watson, S. J., Robinson, T. E., & Akil, H. (2007). Individual differences in the propensity to approach signals vs goals promote different adaptations in the dopamine system of rats. Psychopharmacology, 191(3), 599–607. https://doi.org/10.1007/s00213-006-0535-8

Fraser, K. M., Haight, J. L., Gardner, E. L., & Flagel, S. B. (2016). Examining the Role of Dopamine D2 and D3 Receptors in Pavlovian Conditioned Approach Behaviors. Behavioural Brain Research, 305, 87–99. https://doi.org/10.1016/j.bbr.2016.02.022

Fraser, K. M., & Janak, P. H. (2017). Long-lasting contribution of dopamine in the nucleus accumbens core, but not dorsal lateral striatum, to sign-tracking. European Journal of Neuroscience, 46(4), 2047–2055. https://doi.org/10.1111/ejn.13642

Gheidi, A., Cope, L. M., Fitzpatrick, C. J., Froehlich, B. N., Atkinson, R., Groves, C. K., Barcelo, C. N., & Morrow, J. D. (2020). Effects of the cannabinoid receptor agonist CP-55,940 on incentive salience attribution. Psychopharmacology, 237(9), 2767–2776. https://doi.org/10.1007/s00213-020-05571-3

Hart, A. S., Rutledge, R. B., Glimcher, P. W., & Phillips, P. E. M. (2014). Phasic Dopamine Release in the Rat Nucleus Accumbens Symmetrically Encodes a Reward Prediction Error Term. Journal of Neuroscience, 34(3), 698–704. https://doi.org/10.1523/JNEUROSCI.2489-13.2014

Hearst, E., & Jenkins, H. M. (1974). Sign-tracking: The Stimulus-reinforcer Relation and Directed Action. Psychonomic Society.

Horvitz, J. C., Stewart, T., & Jacobs, B. L. (1997). Burst activity of ventral tegmental dopamine neurons is elicited by sensory stimuli in the awake cat. Brain Research, 759(2), 251–258. https://doi.org/10.1016/S0006-8993(97)00265-5

Jones, S. R., O’Dell, S. J., Marshall, J. F., & Wightman, R. M. (1996). Functional and anatomical evidence for different dopamine dynamics in the core and shell of the nucleus accumbens in slices of rat brain. *Synapse (New York*, N.Y.), 23(3), 224– 231. https://doi.org/10.1002/(SICI)1098-2396(199607)23:3<224::AID-SYN12>3.0.CO;2-Z

Justinova, Z., Munzar, P., Panlilio, L. V., Yasar, S., Redhi, G. H., Tanda, G., & Goldberg, S. R. (2008). Blockade of THC-seeking behavior and relapse in monkeys by the cannabinoid CB(1)-receptor antagonist rimonabant. Neuropsychopharmacology: Official Publication of the American College of Neuropsychopharmacology, 33(12), 2870–2877. https://doi.org/10.1038/npp.2008.21

Kochli, D. E., Keefer, S. E., Gyawali, U., & Calu, D. J. (2020). Basolateral Amygdala to Nucleus Accumbens Communication Differentially Mediates Devaluation Sensitivity of Sign- and Goal-Tracking Rats. Frontiers in Behavioral Neuroscience, 14, 593645. https://doi.org/10.3389/fnbeh.2020.593645

Lefner, M. J., Dejeux, M. I., & Wanat, M. J. (2022). Sex Differences in Behavioral Responding and Dopamine Release during Pavlovian Learning. ENeuro, 9(2). https://doi.org/10.1523/ENEURO.0050-22.2022

Lopez, J. C., Karlsson, R.-M., & O’Donnell, P. (2015). Dopamine D2 Modulation of Sign and Goal Tracking in Rats. Neuropsychopharmacology, 40(9), 2096–2102. https://doi.org/10.1038/npp.2015.68

Lupica, C. R., & Riegel, A. C. (2005). Endocannabinoid release from midbrain dopamine neurons: A potential substrate for cannabinoid receptor antagonist treatment of addiction. Neuropharmacology, 48(8), 1105–1116. https://doi.org/10.1016/j.neuropharm.2005.03.016

Lupica, C. R., Riegel, A. C., & Hoffman, A. F. (2004). Marijuana and cannabinoid regulation of brain reward circuits. British Journal of Pharmacology, 143(2), 227– 234. https://doi.org/10.1038/sj.bjp.0705931

Madayag, A. C., Stringfield, S. J., Reissner, K. J., Boettiger, C. A., & Robinson, D. L. (2017). Sex and Adolescent Ethanol Exposure Influence Pavlovian Conditioned Approach. Alcoholism, Clinical and Experimental Research, 41(4), 846–856. https://doi.org/10.1111/acer.13354

McLaughlin, P. J., Qian, L., Wood, J. T., Wisniecki, A., Winston, K. M., Swezey, L. A., Ishiwari, K., Betz, A. J., Pandarinathan, L., Xu, W., Makriyannis, A., & Salamone, J. D. (2006). Suppression of food intake and food-reinforced behavior produced by the novel CB1 receptor antagonist/inverse agonist AM 1387. Pharmacology, Biochemistry, and Behavior, 83(3), 396–402. https://doi.org/10.1016/j.pbb.2006.02.022

McLaughlin, P. J., Winston, K., Swezey, L., Wisniecki, A., Aberman, J., Tardif, D. J., Betz, A. J., Ishiwari, K., Makriyannis, A., & Salamone, J. D. (2003). The cannabinoid CB1 antagonists SR 141716A and AM 251 suppress food intake and food-reinforced behavior in a variety of tasks in rats. Behavioural Pharmacology, 14(8), 583–588. https://doi.org/10.1097/00008877-200312000-00002

Meyer, P. J., Lovic, V., Saunders, B. T., Yager, L. M., Flagel, S. B., Morrow, J. D., & Robinson, T. E. (2012). Quantifying Individual Variation in the Propensity to Attribute Incentive Salience to Reward Cues. PLOS ONE, 7(6), e38987. https://doi.org/10.1371/journal.pone.0038987

Mikhael, J. G., & Gershman, S. J. (2019). Adapting the flow of time with dopamine. Journal of Neurophysiology, 121(5), 1748–1760. https://doi.org/10.1152/jn.00817.2018

Mirenowicz, J., & Schultz, W. (1994). Importance of unpredictability for reward responses in primate dopamine neurons. Journal of Neurophysiology, 72(2), 1024–1027. https://doi.org/10.1152/jn.1994.72.2.1024

Oleson, E. B., Beckert, M. V., Morra, J. T., Lansink, C. S., Cachope, R., Abdullah, R. A., Loriaux, A. L., Schetters, D., Pattij, T., Roitman, M. F., Lichtman, A. H., & Cheer, J. F. (2012). Endocannabinoids Shape Accumbal Encoding of Cue-Motivated Behavior via CB1 Receptor Activation in the Ventral Tegmentum. Neuron, 73(2), 360–373. https://doi.org/10.1016/j.neuron.2011.11.018

Pitchers, K. K., Flagel, S. B., O’Donnell, E. G., Solberg Woods, L. C., Sarter, M., & Robinson, T. E. (2015). Individual variation in the propensity to attribute incentive salience to a food cue: Influence of sex. Behavioural Brain Research, 278, 462– 469. https://doi.org/10.1016/j.bbr.2014.10.036

Roitman, M. F., Stuber, G. D., Phillips, P. E. M., Wightman, R. M., & Carelli, R. M. (2004). Dopamine operates as a subsecond modulator of food seeking. The Journal of Neuroscience: The Official Journal of the Society for Neuroscience, 24(6), 1265–1271. https://doi.org/10.1523/JNEUROSCI.3823-03.2004

Saddoris, M. P., Cacciapaglia, F., Wightman, R. M., & Carelli, R. M. (2015). Differential Dopamine Release Dynamics in the Nucleus Accumbens Core and Shell Reveal Complementary Signals for Error Prediction and Incentive Motivation. The Journal of Neuroscience: The Official Journal of the Society for Neuroscience, 35(33), 11572–11582. https://doi.org/10.1523/JNEUROSCI.2344-15.2015

Saddoris, M. P., Sugam, J. A., Cacciapaglia, F., & Carelli, R. M. (2013). Rapid dopamine dynamics in the accumbens core and shell: Learning and action. Frontiers in Bioscience (Elite Edition), 5, 273–288. https://doi.org/10.2741/e615

Saddoris, M. P., Wang, X., Sugam, J. A., & Carelli, R. M. (2016). Cocaine Self-Administration Experience Induces Pathological Phasic Accumbens Dopamine Signals and Abnormal Incentive Behaviors in Drug-Abstinent Rats. Journal of Neuroscience, 36(1), 235–250. https://doi.org/10.1523/JNEUROSCI.3468-15.2016

Salamone, J. D., McLaughlin, P. J., Sink, K., Makriyannis, A., & Parker, L. A. (2007). Cannabinoid CB1 receptor inverse agonists and neutral antagonists: Effects on food intake, food-reinforced behavior and food aversions. Physiology & Behavior, 91(4), 383–388. https://doi.org/10.1016/j.physbeh.2007.04.013

Saunders, B. T., & Robinson, T. E. (2012). The role of dopamine in the accumbens core in the expression of Pavlovian-conditioned responses. The European Journal of Neuroscience, 36(4), 2521–2532. https://doi.org/10.1111/j.1460-9568.2012.08217.x

Saunders, B. T., Yager, L. M., & Robinson, T. E. (2013). Cue-evoked cocaine “craving”: Role of dopamine in the accumbens core. The Journal of Neuroscience: The Official Journal of the Society for Neuroscience, 33(35), 13989–14000. https://doi.org/10.1523/JNEUROSCI.0450-13.2013

Schindler, C. W., Redhi, G. H., Vemuri, K., Makriyannis, A., Le Foll, B., Bergman, J., Goldberg, S. R., & Justinova, Z. (2016). Blockade of Nicotine and Cannabinoid Reinforcement and Relapse by a Cannabinoid CB1-Receptor Neutral Antagonist

AM4113 and Inverse Agonist Rimonabant in Squirrel Monkeys. Neuropsychopharmacology: Official Publication of the American College of Neuropsychopharmacology, 41(9), 2283–2293. https://doi.org/10.1038/npp.2016.27

Schultz, W., Dayan, P., & Montague, P. R. (1997). A neural substrate of prediction and reward. *Science (New York*, N.Y.), 275(5306), 1593–1599. https://doi.org/10.1126/science.275.5306.1593

Stelly, C. E., Girven, K. S., Lefner, M. J., Fonzi, K. M., & Wanat, M. J. (2021). Dopamine release and its control over early Pavlovian learning differs between the NAc core and medial NAc shell. Neuropsychopharmacology: Official Publication of the American College of Neuropsychopharmacology, 46(10), 1780–1787. https://doi.org/10.1038/s41386-020-00941-z

Tomie, A. (1996). Locating reward cue at response manipulandum (CAM) induces symptoms of drug abuse. Neuroscience and Biobehavioral Reviews, 20(3), 505– 535. https://doi.org/10.1016/0149-7634(95)00023-2

Valyear, M. D., Glovaci, I., Zaari, A., Lahlou, S., Trujillo-Pisanty, I., Andrew Chapman, C., & Chaudhri, N. (2020). Dissociable mesolimbic dopamine circuits control responding triggered by alcohol-predictive discrete cues and contexts. Nature Communications, 11(1), 3764. https://doi.org/10.1038/s41467-020-17543-4

Ward, S. J., & Dykstra, L. A. (2005). The role of CB1 receptors in sweet versus fat reinforcement: Effect of CB1 receptor deletion, CB1 receptor antagonism (SR141716A) and CB1 receptor agonism (CP-55940). Behavioural Pharmacology, 16(5–6), 381–388. https://doi.org/10.1097/00008877-200509000-00010

Ward, S. J., Walker, E. A., & Dykstra, L. A. (2007). Effect of cannabinoid CB1 receptor antagonist SR141716A and CB1 receptor knockout on cue-induced reinstatement of Ensure and corn-oil seeking in mice. Neuropsychopharmacology: Official Publication of the American College of Neuropsychopharmacology, 32(12), 2592–2600. https://doi.org/10.1038/sj.npp.1301384

Wenzel, J. M., Oleson, E. B., Gove, W. N., Cole, A. B., Gyawali, U., Dantrassy, H. M., Bluett, R. J., Dryanovski, D. I., Stuber, G. D., Deisseroth, K., Mathur, B. N., Patel, S., Lupica, C. R., & Cheer, J. F. (2018). Phasic Dopamine Signals in the Nucleus Accumbens that Cause Active Avoidance Require Endocannabinoid Mobilization in the Midbrain. Current Biology, 28(9), 1392–1404.e5. https://doi.org/10.1016/j.cub.2018.03.037

West, E. A., & Carelli, R. M. (2016). Nucleus Accumbens Core and Shell Differentially Encode Reward-Associated Cues after Reinforcer Devaluation. Journal of Neuroscience, 36(4), 1128–1139. https://doi.org/10.1523/JNEUROSCI.2976-15.2016

Zahm, D. S. (1999). Functional-anatomical implications of the nucleus accumbens core and shell subterritories. Annals of the New York Academy of Sciences, 877, 113– 128. https://doi.org/10.1111/j.1749-6632.1999.tb09264.x

